# Addressing Enzymatic-Independent Tumor-Promoting Function of NAMPT via PROTAC-Mediated Degradation

**DOI:** 10.1101/2021.09.13.460066

**Authors:** Xiaotong Zhu, Haixia Liu, Li Chen, Yong Cang, Biao Jiang, Xiaobao Yang, Gaofeng Fan

**Affiliations:** School of Life Science and Technology, ShanghaiTech University, Shanghai 201210, China; School of Physical Science and Technology, ShanghaiTech University, Shanghai 201210, China; Shanghai Institute for Advanced Immunochemical Studies, ShanghaiTech University, Shanghai 201210, China; University of Chinese Academy of Sciences, Beijing 100049, China; CAS Key Laboratory of Synthetic Chemistry of Natural Substances, Shanghai Institute of Organic Chemistry, Chinese Academy of Sciences, Shanghai 200032, China; Gluetacs Therapeutics (Shanghai) Co., Ltd., Zhangjiang Hi-Tech Park, Shanghai, 201210, China

**Keywords:** PROTAC, iNAMPT, eNAMPT, enzymatic-independent, NAD^+^

## Abstract

The rate-limiting enzyme of salvage pathway for NAD^+^ synthesis, NAMPT, is aberrantly overexpressed in a variety of tumor cells and is a poor prognosis factor for patient survival. NAMPT plays a major role in tumor cell proliferation, acting concurrently as an NAD^+^ synthase and unexpectedly, an extracellular ligand for several tumor-promoting signaling pathways. While previous efforts to modulate NAMPT activity were limited to enzymatic inhibitors with low success in clinical studies, protein degradation offers a possibility to simultaneously disrupt NAMPT’s enzyme activity and ligand capabilities. Here, we report the development of two highly selective NAMPT-targeted proteolysis-targeting chimeras (PROTACs), which promoted rapid and potent NAMPT degradation in a cereblon-dependent manner in multiple tumor cell lines. Notably, both PROTAC degraders outperform a clinical candidate, FK866, in killing effect on hematological tumor cells. These results emphasize the importance and feasibility of applying PROTACs as a better strategy for targeting proteins like NAMPT with dual tumor-promoting functions, which are not easily achieved by conventional enzymatic inhibitors.

## Introduction

As a new format of drug design that is different from traditional small molecule inhibitors, PROTAC is composed of three key parts: target protein binder, linker and E3 ubiquitin ligase binder.^1^ Since the first small molecule PROTAC was reported in 2001 by Prof. Craig Crews at Yale University, the field has grown rapidly. Over the past 20 years, thousands of PROTAC molecules have been developed by the scientific and industrial community. At present, PROTAC molecules targeting the androgen receptor (AR), estrogen receptor (ER), bruton tyrosine kinase (BTK), bromodomain-containing protein 9 (BRD9), B-cell lymphoma extra large (Bcl-X_L_) and interleukin-1 receptor-associated kinase 4 (IRAK4) have entered the clinical development stage.^2 3^

As an important small molecule metabolite, nicotinamide adenine dinucleotide (NAD^+^) is widely involved in a series of biochemical reactions in cell energy metabolism, such as glycolysis, oxidative phosphorylation and fatty acid oxidation. It also plays a versatile role in signaling transduction events as a co-substrate, including DNA damage repair and protein deacetylation. Nicotinamide phosphoribosyltransferase (NAMPT) is the rate-limiting enzyme that catalyzes the synthesis of nicotinamide mononucleotide (NMN) from nicotinamide (NAM) in the mammalian NAD^+^ synthetic salvage pathway.^4^ There are two forms of NAMPT that exist, including intracellular NAMPT (iNAMPT) and extracellular NAMPT (eNAMPT). A body of literature has demonstrated a critical role of NAMPT in promoting tumor cell proliferation and dedifferentiation.^5–8^ Due to increased NAD^+^ and ATP catabolism, tumor cells are more sensitive to iNAMPT inhibition than normal cells.^9–10^ Interestingly, eNAMPT can be released and acts as an oncogenic cytokine on a variety of cells, with TLR4 as its putative receptor.^11–12^ Elevation of eNAMPT in the plasma of patients has been documented in a variety of human malignancies, including astrocytoma, myeloma, and male oral squamous cells, gastric cancer, endometrial cancer, hepatocellular carcinoma, colorectal cancer, and aggressive breast cancer.^13–21^ Therefore, it is crucial to antagonize functions of both forms of NAMPT for better treatment of related tumor diseases, and apparently traditional small molecule inhibitor comes a little bit short for covering both.

Research led by a team in the UK has recently proposed a systematic approach to assess the PROTAC ability of protein targets.^2^ Using this approach, they identified 1067 PROTAC targets that have not yet been reported in the literature, providing potential new opportunities for PROTAC-based drug development. Four major reasons ensure NAMPT as one of the leading candidates among 1067 uncharacterized targets under the “Discovery Opportunity” category: (1) Cellular location of the protein fits the optimal efficacy for PROTAC molecule; (2) The half-lives of 9699 proteins have been determined by using SILAC (stable isotope labeling by amino acids in cell culture) technology^22^, which ranges from 10 to >1000 h. The half-life of NAMPT protein across 5 different cell types is within the range of 45~153 hrs^22^, suitable for protein degradation-based inhibitor design; (3) 24 potential ubiquitination sites in NAMPT has been revealed^23^; (4) Multiple small-molecule ligands for NAMPT have been deposited in ChEMBL^24^, all with a promising binding affinity to further pursue PROTAC molecule design. Of note, the pChEMBL value (the negative logarithm of the reported activity such as IC_50_, EC_50_ or *K*_d_) of NAMPT inhibitor FK866 is 10.52.

Here, we report the development of two highly selective NAMPT-targeted PROTACs, which promoted rapid and potent degradation of both iNAMPT and eNAMPT in a cereblon-dependent manner in multiple tumor cell lines. Notably, both PROTAC degraders outperform their original target binder, FK866, in killing effect on hematological tumor cells. These results highlight the tremendous advantage of applying PROTACs to target proteins like NAMPT with dual tumor-promoting functions, which are not easily achieved by conventional enzymatic inhibitors.

## Experimental Methods

### PROTAC Synthesis

Detailed information regarding synthesis procedure and NMR analytical data could be found in the supplementary materials.

### Cell culture

HEK293T (CRL-3249, ATCC), SW620 (CCL-227, ATCC), HT29 (HTB-38, ATCC), MCF-7 (HTB-22, ATCC), MOLT4 (CRL-1582, ATCC), Jurkat (TIB-152, ATCC), HL60 (CCL-240, ATCC), BEL7404 were used for cellular assay. HEK293T, SW620, HT29, MCF-7, BEL7404 cells were grown in DMEM supplemented with 10% FBS and Pen/Strep at 37 °C with 5% CO_2_. Jurkat, MOLT4 and HL60 cells were grown in RPMI 1640 medium supplemented with 10% FBS and Pen/Strep.

### Establishment of NAMPT fluorescence reporter cell line

The procedures used in Yao, X. et al^25^ were followed to generate the NAMPT fluorescence reporter cell line. In brief, first, the left and right homology arms (800 bp HAL and 800bp HAR) flanking the stop codon of the *NAMPT* gene were amplified using HEK293T cell genomic DNA as a template. The homologous recombination method was applied to obtain the final linearized donor (800bp HAL-GGSG-GFP-3xFLAG-800bp HAR).

Next, use the online CRISPR tool (http://chopchop.cbu.uib.no/) to design sgRNAs for the target region (flanking the stop codon of the *NAMPT* gene). Multiple sgRNAs were ligated to the linearized PX330 vector (Addgene, #42230). To evaluate the targeting efficiency of different sgRNAs, transiently transfect the corresponding pX330 vector into HEK293T cells with lipofectamine3000 for 48 hrs. Extract the genomic DNA, and sequence the sgRNA-targeted site to calculate the editing efficiency by using the online tool https://tide.nki.nl/. The 23nt NAMPT-sgRNA target sequence (ACATCATTAGGCTTTATGACTGG) was finally selected.

Lastly, HEK293T cells were transfected with total of 12 μg plasmids (sgRNA PX330: donor = 1:3) for 96 hours. GFP-positive cells were then sorted into 96-well plates for single clone cell line establishment. The insertion of GFP fluorescence reporter was verified by genomic DNA sequencing, western blot, fluorescence microscope imaging and flow cytometry (FACS) analysis.

### Flow cytometry

HEK293T^GFP-NAMPT^ cells were seeded in a 6-well plate at 1×10^6^ cells/well and exposed to different concentrations of **630120** or **630121**. After 24 hrs, cells were harvested and washed by PBS 3 times. The protein level of NAMPT was then indicated by the value of GFP fluorescence, which could be measured by flow cytometry (BD LSRFortessa).

### NAD^+^ concentration measurement

HL60 and MOLT4 cells (1 × 10^6^ cells/sample) were harvested and intracellular NAD^+^ levels were determined by using an NAD^+^/NADH assay kit (Beyotime, cat. no. S0175) according to the manufacturer’s instructions.

### Enrichment of secreted protein(s) in the culture supernatant

HL60 and MOLT4 cells were incubated in a 6-well plate with 2 ml RPMI 1640 full medium for 24 and 48 hrs, respectively. Then, the supernatant medium was collected and added to the Amicon® Ultra-15 Centrifugal Filter (Millipore, cat. no. UFC901096) for secreted protein enrichment according to the manufacturer’s instructions.

### Immunoblotting analysis

Cell extracts were prepared in 1% NP40 lysis buffer (20 mM Hepes pH 7.5, 150 mM NaCl, 1% NP40, 50 mM NaF, 1 mM Na_3_VO_4_, 10% glycerol, and protease inhibitor cocktail from Roche) at 4 °C for 15 mins. Total protein concentration was determined with the BCA kit (Beyotime, cat. no. P0010). An equal amount of proteins was subjected to SDS-PAGE electrophoresis, followed by nitrocellulose membrane transfer at 95 Volts and 4 °C for 90 mins. After the transfer, the membrane was blocked with 5% nonfat milk in TBST for 1 hr at room temperature. Incubate the membrane with the primary antibody at 4 °C overnight. The membrane was then washed 3 times with TBST at room temperature, and incubated with secondary antibody conjugated with horseradish peroxidase (Cell Signaling Technology) at room temperature for 1 hr. After the final wash with TBST for 3 times, the membrane was incubated with Super Signal West Pico PLUS Chemiluminescent Substrate (Thermo Fisher) and developed on GE developer (AI680UV).

The following antibodies were purchased from the indicated suppliers and used for immunoblotting analysis: anti-GAPDH (Abclonal, cat. no. A19056); anti-NAMPT (Proteintech, cat. no. 66385-1-lg); Anti-Phospho-Erk1/2 (Cell signaling technology, cat. no. 4370); anti-Erk1/2 (Cell signaling technology, cat. no. 4695).

### Silver Staining

Protein samples were first subjected to SDS-PAGE electrophoresis, followed by silver staining using a Fast Silver Stain Kit (Beyotime, cat. no. P0017S) according to the manufacturer’s instructions.

### CTG assay

Cell Titer-Glo (CTG) luminescent cell viability assay (Promega) was used to calculate the IC_50_ of PROTAC compounds. In brief, PROTAC compounds, diluted tenfold from a concentration of 10 μM to a final concentration of 0.01 nM, were incubated with pre-seeded cells in 96-well plates for an indicated time interval. Then CTG reagent was added to each well and mixed for 10 mins on an orbital shaker, followed by luminescence measurement with SpectraMax i3 (MD).

### *In vitro* killing assay

FK866 or PROTAC compounds treated cells were stained with DAPI (1:250), followed by flow cytometry analysis. We used the ratio of DAPI positive to calculate the killing efficacy of these compounds.

### Quantitative proteomic analysis

HCT116 cells, treated with either DMSO or PROTAC compounds (10 nM) for 24 hrs, were lysed in buffer containing 50 mM NH_4_HCO_3_, 8 M urea and protease inhibitor (Roche, 05892791001) with sonication on ice. The final protein concentration was determined using a BCA kit (Beyotime, cat. no. P0012S). The lysate was then incubated with 5 mM 1,4-dithiothreitol (DTT) at 37 °C for 1 hr, followed by 10 mM 2-Iodoacetamide (IAM) treatment at room temperature for 45 mins. Next, add 50 mM NH_4_HCO_3_ into the protein solution, followed by adding sequencing grade Trypsin at the ratio of 1:50 (Promega, cat. no. V5111) for digestion at 37 °C overnight. On the next day, 10% trifluoroacetic acid (TFA) was added to the protein digest to a final concentration of 0.4%, followed by desalting the peptide by Titansphere Phos-TiO kit according to the manufacturers instructions (GL Sciences, cat. no. 5010-21312). The peptide was vacuum-dried and stored at −20 °C or −80 °C. The peptide segment was analyzed and processed by the instrument (Thermo Fisher, Q Exactive HF-X). Two biological replicates were performed for each sample. The **630120**/DMSO and **630121**/DMSO ratios represent the effects of **630120** and **630121**. The scatter plot was drawn using GraphPad Prism 9.

## Results

### Degradation of endogenous NAMPT proteins is mainly through lysosome rather than proteasome

Target gene with fluorescence reporter will not only benefit basic research regarding the regulation of transcriptional activation and post-translation modification but also facilitate translational attempts in screening and identifying small molecule inhibitors like PROTAC or molecular glue in a high-throughput manner. For these purposes, we inserted a GFP-FLAG tag just before the stop codon of the *NAMPT* gene in HEK293 T cells (Figure 1A) by adopting Tild-CRISPR (targeted integration with linearized dsDNA-CRISPR) strategy ^25–28^. After FACS sorting, single clone recovering and biochemical validation, we successfully established one reporter line for NAMPT. Immunoblotting analysis with cell lysates illustrated two major bands recognized by NAMPT antibody, one was ~50 kDa which represents endogenous NAMPT and the other was ~75kDa which represents GFP-FLAG-fused gene (Figure 1B). Precise genome integration by Tild-CRISPR strategy was further confirmed by genomic DNA sequencing, western blot, fluorescence microscope imaging and flow cytometry (FACS) analysis (Figure S1A-B). Notably, we observed a remarkable fluorescence intensity decrease upon shRNA-mediated NAMPT gene knock-down in this reporter cell line (Figure 1C), in conjunction with the loss of NAMPT protein expression (Figure 1D), demonstrating the on-target GFP-FLAG tag insertion within NAMPT open reading frame (ORF).

**Figure 1.**
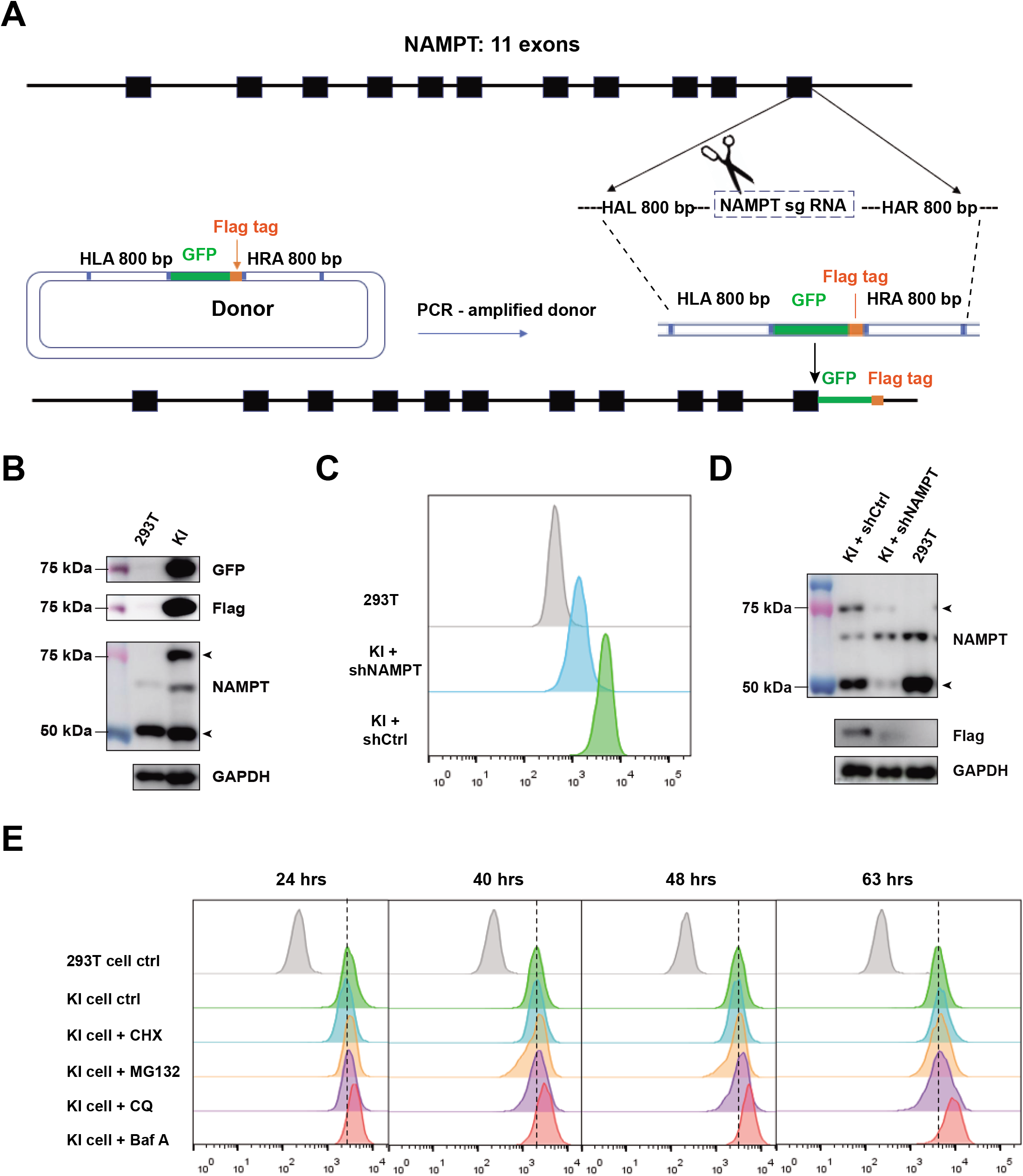
Degradation of endogenous NAMPT proteins is mainly through lysosome rather than proteasome. (A) Schematic overview of construction of NAMPT knock-in HEK293T cell line using Tild-CRISPR. HLA, left homology arm; HRA, right homology arm. (B) Western blot verification of NAMPT protein before and after GFP-FLAG tag insertion. (C and D) Flow cytometry analysis of GFP fluorescence intensity (C) and Western blot verification of NAMPT protein expression (D) in parental HEK293T cells and GFP-FLAG knock-in cells with control or NAMPT shRNA infection. (E) Flow cytometry analysis of GFP fluorescence intensity in HEK293T cells and GFP-FLAG knock-in cells after treating with CHX, MG132, CQ and Baf A for 24h, 40h, 48h and 63h.

Given the important role of NAMPT in NAD^+^ synthesis, we first utilized this fluorescence reporter cell line to investigate the regulatory mechanism of its protein stability. Four reagents were applied in our study, including cycloheximide (CHX) - a protein synthesis inhibitor, MG132 - a proteasome inhibitor, chloroquine (CQ) - an autophagy inhibitor by destroying the structure and function of lysosome and bafilomycin A1 (BafA) - a lysosome maturation inhibitor by inhibiting the H^+^-ATPase function. CHX treatment revealed that NAMPT protein was quite stable, with a half-life of more than 60 hrs. Meanwhile, neither MG132 nor CQ could accumulate NAMPT protein (Figure 1E), indicating both proteasome and autophagy machineries make a limited contribution to the turnover of this protein. However, BafA treatment overtly enhanced the GFP fluorescence signal (Figure 1E), suggesting degradation of NAMPT protein is mainly through the lysosome pathway.

### Design and Synthesis of NAMPT-targeting PROTACS

Though NAMPT protein was found to be degraded mainly through lysosome rather than proteasome, the facts that NAMPT is one of the “Discovery Opportunity” candidates^2^ and the necessity of targeting both iNAMPT and eNAMPT in treating cancers strongly prompted us to explore the feasibility of PROTAC strategy. NAMPT specific inhibitor FK866 and lenalidomide were selected as degradation target binder and E3 ligase cereblon binder, respectively. After rounds of optimization, we adopted piperazine as an extended group and 5 or 6 atoms alkyl-chain as a linker for **SIAIS630120** and **SIAIS630121** (Fig. 2A-B), respectively. The whole scheme for the design and synthesis of these two NAMPT-targeting PROTACs was illustrated in Scheme 1 and details in Supporting Information.

**Figure 2.**
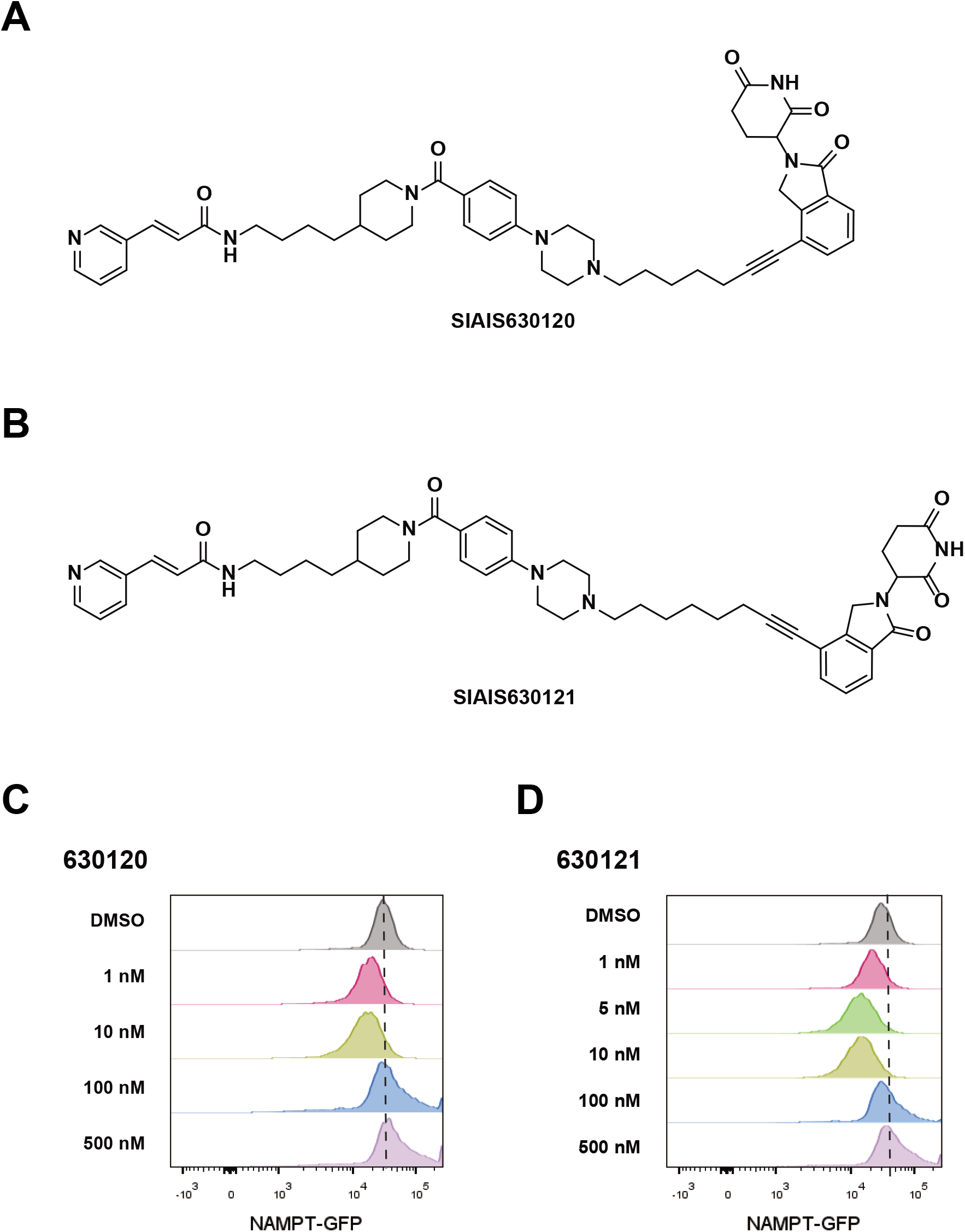
The design and synthesis of NAMPT-targeting PROTAC compounds and t heir efficacy *in vitro*. (A-B) Chemical structures of **630120** (A) and **630121** (B). (C-D) Degradation efficacy evaluation of PROTAC compounds **630120** (C) and **630121** (D) after 24-hr incubation with HEK293T^NAMPT-GFP^ cells. The protein level of NAMPT was indicated by the intensity of GFP fluorescence measured by flow cytometry (n = 3).

**Scheme 1.**
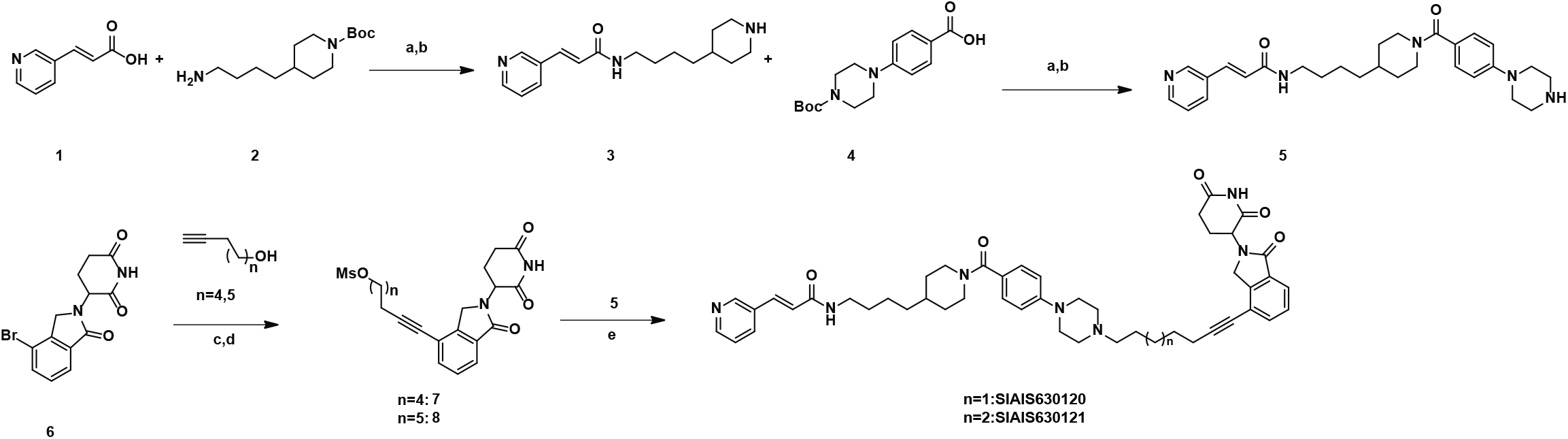
Synthetic routes for NAMPT PROTAC degraders SIAIS630120 and SIAIS630121. Reagents and conditions: (a) HATU, DIPEA, DMF, 60 °C, 12hrs; (b) TFA, DCM, rt, 12hrs; (c) Pd(PPh_3_)_2_Cl_2_, CuI, DMF, TEA, 80 °C, 8hrs; (d) MsCl, TEA, DCM,15 mins; (e) DIPEA, DMF, 60 °C, 6hrs.

Next, we evaluated the efficacy of these two PROTAC molecules in degrading NAMPT protein in HEK293T^NAMPT-GFP^ cells, using fluorescence intensity as readout by flow cytometry analysis. NAMPT-GFP was retained in those cells treated with DMSO but diminished in cells treated with **SIAIS630120** or **SIAIS630121** at relatively low concentrations (Fig. 3C-D). This fluorescence intensity loss was significantly reduced at a higher concentration of treatment, probably due to the hook effect of PROTAC molecules.^29^ In sum, we successfully designed two NAMPT-oriented PROTAC compounds, **SIAIS630120** (hereinafter referred to as **630120**) and **SIAIS630121** (hereinafter referred to as **630121**), with effective degradation of the target protein.

**Figure 3.**
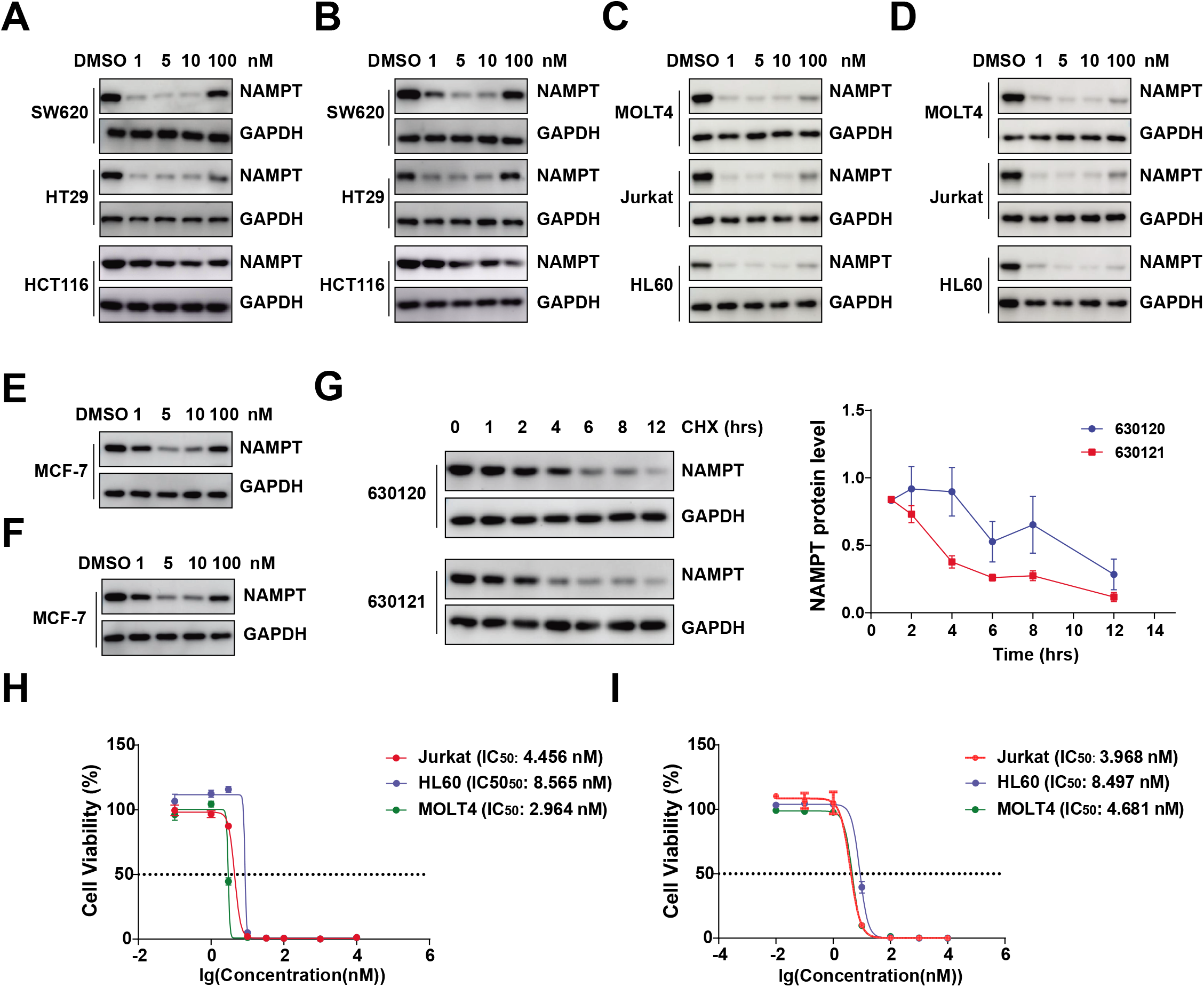
Functional characterization of 630120 and 630121 in multiple tumor cell lines. (A-B) NAMPT degradation assay with **630120** or **630121** treatment in CRC cell lines. SW620, HT29 and HCT116 cells were treated with DMSO, **630120** (A) or **630121** (B) for 24 hrs at the indicated concentrations, followed by Western blot. (C-D) NAMPT degradation assay with **630120** or **630121** treatment in hematological tumor cell lines. MOLT4, Jurkat and HT29 cells were treated with DMSO, **630120** (C) or **630121** (D) for 24 hrs at the indicated concentrations, followed by Western blot. (E-F) NAMPT degradation assay with **630120** or **630121** treatment in breast cancer cell line. MCF-7 cells were treated with DMSO, **630120** (E) or **630121** (F) for 24 hrs at the indicated concentrations, followed by Western blot. (G) Degradation kinetics of NAMPT by two PROTAC compounds in Jurkat cells. Jurkat cells were treated with 10nM **630120** or **630121**. At the indicated time intervals, cell lysates were extracted for Western blot with NAMPT antibody. NAMPT bands were quantified with ImageJ. Data were shown as mean ± SD. Each sample group comprised three replicates. (H-I) CTG luminescent cell viability assay (Details in Methods) was performed to calculate IC_50_ of **630120** (H) and **630121** (I), respectively in Jurkat, HL60 and MOLT4 cell lines (96-hour treatment). Data were shown as mean ± SD. Each sample group comprised three replicates.

### Functional characterization of 630120 and 630121 in multiple tumor cell lines

NAMPT is aberrantly upregulated in colorectal cancer (CRC), breast cancer, and hematological malignancies as well, and is correlated with advanced metastasis and poor prognosis^5^. Genetic or pharmacological inhibition of NAMPT significantly reduced the growth of tumor cells in multiple subtypes and promote apoptosis both *in vitro* and *in vivo*.^30–32^ Therefore, we attempted to assess the anti-tumor effect of our NAMPT-targeting PROTAC compounds in a panel of tumor cell lines, starting with their protein degradation properties.

Firstly, these two PROTAC compounds illustrated excellent compacity in degrading the target protein NAMPT. As shown in Fig 3, both **630120** and **630121** showed overtly NAMPT protein degradation in three colorectal cancer cell lines SW620, HT29 and HCT116 (Fig. 3A-B). In addition, effective degradation of NAMPT proteins was also observed in hematological tumor cell lines, including MOLT4, Jurkat and HL60 cells (Fig 3C-D), as well as cell lines of other types of solid tumors, such as breast cancer cell line MCF-7 (Fig 3E-F).

Secondly, these two PROTAC compounds elicited the degradation effect of NAMPT protein in a very rapid manner. As shown in Fig. 3G, it took 4 hours for 10nM **630121** and 6 hours for 10nM **630120** to degrade 50% of endogenous NAMPT protein in Jurkat cells.

Thirdly, both PROTAC compounds exhibited high potency at very low concentrations. Cell Titer-Glo (CTG) luminescent cell viability assay^33^ was applied to calculate the IC_50_ values of **630120** and **630121**. As shown in Fig. 3H, the IC_50_ values of **630120** in Jurkat, HL60 and MOLT4 cell lines were 4.456 nM, 8.565 nM and 2.964 nM; the corresponding value for **630121** was 3.968 nM, 8.497 nM and 4.681 nM (Fig. 3I).

Lastly, we evaluated the pharmacokinetic (PK) of **630120** and **630121** in Sprague Dawley rats. AUC (area under the plasma concentration-time curve) values of **630120** and **630121** were 735.91 and 452.49 hr*ng/mL, respectively, after dosing 2 mg/kg intravenously. Furthermore, both **630120** and **630121** possessed oral availability with 13.1% and 23.2% post-dosing 10 mg/kg in rats (Figure S2A-B). Compare with **630120**, **630121** illustrated better pharmacokinetic profiles in oral availability, with the highest plasma concentration of a single 10 mg/kg p.o. dose around 134.86 ng/ml.

### 630120 and 630121 specifically degrade NAMPT protein in a cereblon and proteasome-dependent manner

To further evaluate the degradation selectivity of both **630120** and **630121**, we performed a mass spectrometry-based quantitative proteomic analysis in the HCT116 CRC cell line in the absence and presence of the NAMPT-targeting PROTAC compound. Compared to DMSO-treated control samples, **630120** and **630121**-treated cells showed 78.1% (p=8.06*10^-4^) and 86.2% (p=4.72*10^-3^), respectively, in reduction of NAMPT protein (Figure 4A-B). Most importantly, both compounds exhibited minimal off-target side effects to any other proteins detected by MS (Figure 4A-B, no hits within p<0.01 and log_2_FC<-1 area), indicating the ideal selectivity of these two PROTAC degraders.

**Figure 4.**
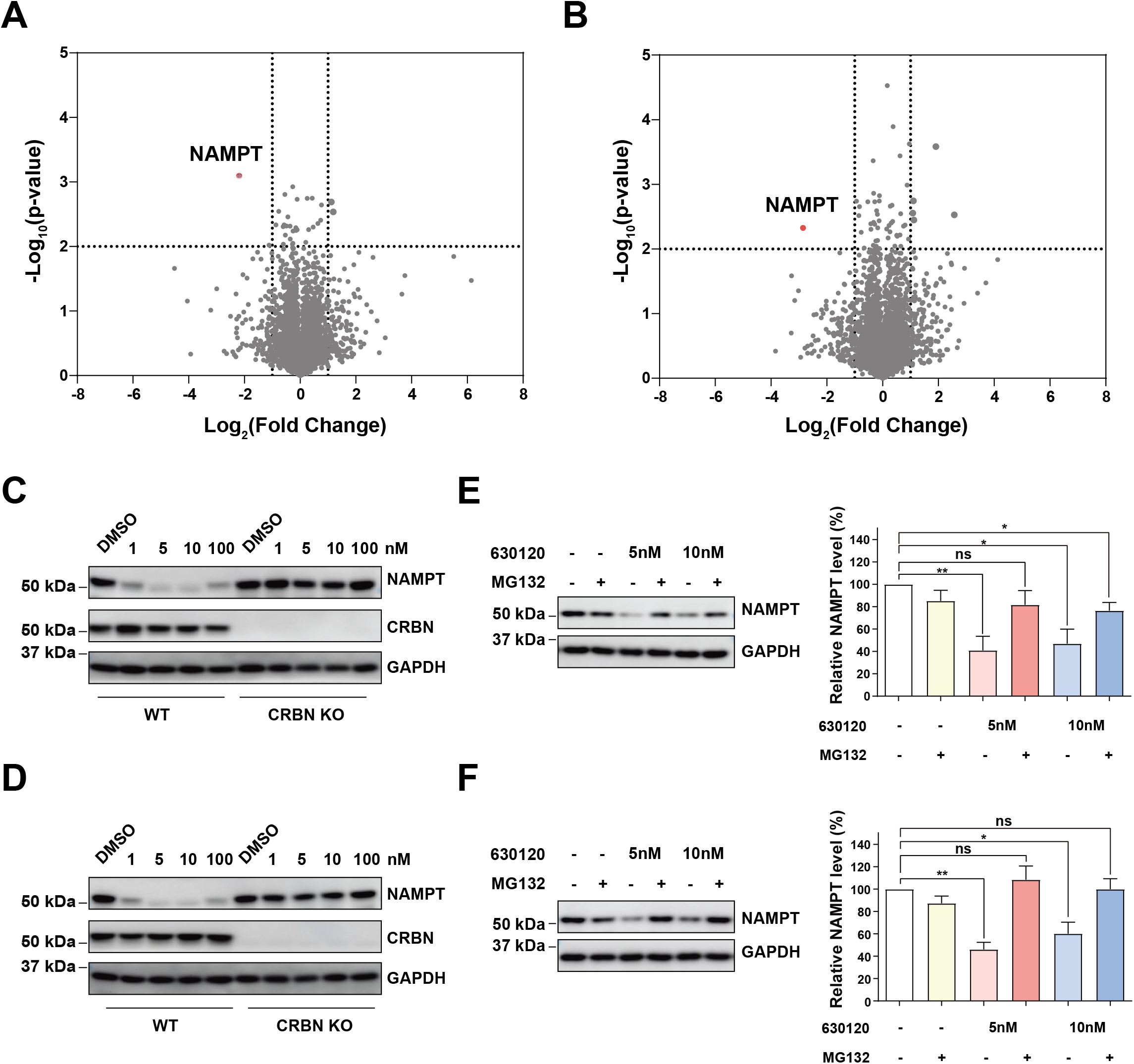
630120 and 630121 specifically degrade NAMPT protein in a cereblonand proteasome-dependent manner. (A-B) Mass spectrometry-based quantitative proteomic analysis in HCT116 CRC cell line in the absence (DMSO) and presence of the NAMPT-targeting PROTAC compound **630120** (A,10 nM, 24 hrs) and **630121** (B,10 nM, 24 hrs). Each experiment had two biological replicates. (C-D) NAMPT degradation induced by **630120** or **630121** is E3 ligase cereblon-dependent. Liver cancer cell lines BEL7404^WT^ and BEL7404^CRBN-KO^ were treated with DMSO, **630120** (C) and **630121** (D) for 24 hrs at the indicated concentrations, followed by Western blot. (E-F) NAMPT degradation induced by **630120** or **630121** is proteasome-dependent. HCT116 cells were pre-treated with proteasome inhibitor MG132 (10 μM) for 2 hrs, followed by **630120** (E) or **630121** (F) treatment for another 8 hrs. Cell lysates were extracted for Western blot with NAMPT antibody. GAPDH was also probed as a loading control. NAMPT bands were quantified with ImageJ. Data were shown as mean ± SD. Each sample group comprised three replicates.

The Ubiquitin-Proteasome System (UPS) is the primary intracellular pathway for cells to clear the damaged or those no longer required proteins. The PROTAC technology allows the UPS system to be chemically co-opted and aimed to degrade a specific target protein.^34^ Next, we attempted to determine whether the **630120** or **630121**-induced NAMPT degradation is dependent on the ubiquitin-proteasome system, even though endogenous NAMPT protein undergoes lysosome-mediated degradation *per se* (Fig. 1). As shown in Fig. 4C and 4D, both **630120** and **630121** triggered robust degradation of NAMPT in parental liver cancer cell line BEL7404; however, CRISPR-Cas9 meditated E3 ligase cereblon knockout significantly blocked the turnover of the NAMPT protein, indicating cereblon is required for PROTAC compounds induced NAMPT degradation. Furthermore, the addition of proteasome inhibitor MG132 also effectively prevented the degradation of NAMPT protein initiated by these two compounds (Fig. 4E-F), confirming the necessity of proteasome machinery in NAMPT clearance.

### NAMPT-targeting PROTAC compounds outperformed FK866 in killing effect on hematological tumor cells by degrading eNAMPT

NAMPT is a rate-limiting enzyme that catalyzes the synthesis of nicotinamide adenine dinucleotide NAD^+^ through the salvage pathway. Our previous work has demonstrated that hematologic cells are more rely on this pathway for NAD^+^ synthesis.^35^ Once blocked, there will be a profound reduction in intracellular concentration of NAD^+^. So next, we want to examine the change of NAD^+^ levels upon treatment with **630120** or **630121**, in parallel comparison with the enzymatic inhibitor of NAMPT-FK866. As shown in Fig. 5A, the intracellular NAD^+^ level of the HL60 cell line after 18-hr exposure to 10nM FK866, 10nM **630120** or 10nM **630121**, respectively, was severely dropped to a similar level, which was ~80% depletion compared to the untreated group. The same results were also observed in another hematologic tumor cell line MLOT4, with approximately 70% reduction (Fig. 5B). Hence, these two PROTAC compounds were equivalently functional as FK866 in terms of NAD^+^ synthesis activity blockade of NAMPT.

**Figure 5.**
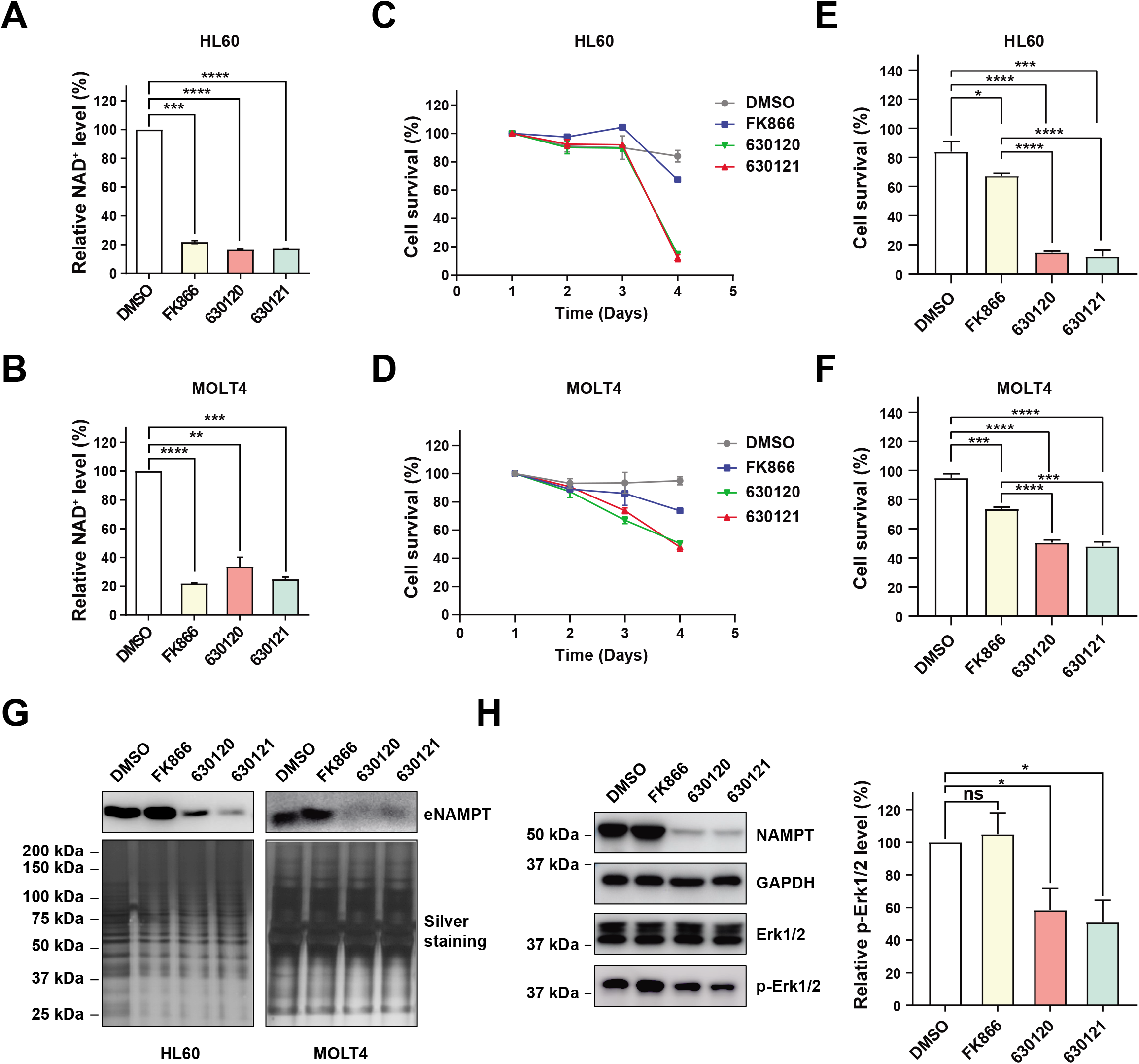
NAMPT-targeting PROTAC compounds outperformed FK866 in killing effect on hematological tumor cells by degrading eNAMPT. (A-B) HL60 (A) and MOLT4 (B) cell lines were treated with 10nM FK866, 10nM **630120** or 10nM **630121**, respectively, for 18 hrs, followed by quantitative analysis of intracellular NAD^+^ level (Details in **Methods**). Data were shown as mean ± SD. Each sample group comprised three replicates. ** p < 0.01; *** p < 0.001; **** p < 0.0001. (C-D) *In vitro* killing assay. HL60 (C) and MOLT4 (D) cell lines were treated with 10nM FK866, 10nM **630120** or 10nM **630121**, respectively, for consecutive 4 days. The cell survival ratio was examined by DAPI-staining based flow cytometry. Data were shown as mean ± SD. Each sample group comprised three replicates. Statistics on the fourth day after incubation (right).* p < 0.05; *** p < 0.001;**** p < 0.0001. (E) eNAMPT level analysis in cell culture supernatants from HL60 and MOLT4. HL60 and MOLT4 cells were treated as indicated (10 nM concentration) for 24 hrs and 48 hrs, respectively. The cell culture supernatant was collected for immunoblotting analysis against eNAMPT. (F) Inhibitory effects of **630120** and **630121** on Erk1/2 signaling. HL60 cells were treated with DMSO, 10nM FK866,10nM **630120** or 10nM **630121**, respectively, for 24 hrs. Cell extracts were collected and subjected to Western blot with antibodies against phosphor- and total Erk1/2. Total protein was probed as a loading control. Phosphor-Erk1/2 bands were quantified with ImageJ. Data were shown as mean ± SD. Each sample group comprised three replicates.

Given the high dependence on NAMPT-mediated NAD^+^ production, we further assessed the fitness of these hematologic tumor cells upon treatment with **630120** or **630121**, in comparison again with FK866. Interestingly, both PROTAC compounds achieved ~95% killing effect on HL60 cells at Day 4, whereas less than 40% cell death was observed in the FK866-treated group (Fig. 5C). Although both PROTAC compounds encountered more tolerance with MOLT4, exhibiting ~50% killing effect at Day 4, it was still 20% better than FK866 (Fig. 5D). Therefore, these two PROTAC compounds outperformed FK866 in killing effect on hematological tumor cells.

NAMPT exists in two forms in mammals: intracellular NAMPT(iNAMPT) in cytoplasm and nucleus and extracellular NAMPT(eNAMPT) in plasma or extracellular space.^36^ eNAMPT behaves like a cytokine by recognizing and activation its putative receptor Toll-like receptor 4 (TLR4).^37^ A body of evidence supports a versatile role of eNAMPT in a different stage of tumor development, including primary tumor cell proliferation^38^, anti-apoptosis^15^,^39^ and epithelial-mesenchymal transformation (EMT)^40^. We observed a more potent killing effect of hematologic tumor cell lines with NAMPT-targeting PROTAC compounds over intracellular enzymatic inhibitor FK866, and we reason this outperformance was due to simultaneous blockade of both forms of NAMPT.

To test this hypothesis, we collected culture supernatants from FK866 or PROTAC compounds treated HL60 and MOLT4 cells, and assess the level of secreted eNAMPT by immunoblotting analysis. We witnessed a significant decrease of eNAMPT in both samples treated with PROTAC compounds, and this decrease was observed in both hematologic tumor cell lines (Fig. 5E). Given lack of the evidence to support the appearance of the UPS system outside the cell, we believe this diminished expression was due to the depletion of iNAMPT by PROTAC, which in turn severely downregulated the secretion of eNAMPT. Interestingly, instead of decreased, FK866 treatment even slightly increased the expression of eNAMPT in both cell lines (Fig. 5E). This was probably due to feedback regulation of cells to respond to FK866 insult and to maintain a minimal concentration of NAD^+^ for survival.

To further understand the additional mechanistic impact of these two PROTAC compounds, we investigated the change of downstream signaling pathways of cells treated with either FK866 or NAMPT degraders. Compared to FK866-treated or the control group, a substantial decrease in MAPK kinase Erk1/2 activation was observed in both PROTAC compounds treated groups (Fig. 5F), which may explain the superior anti-tumor survival and proliferation effects of these molecules seen previously.

## Discussion

NAD exists in two forms: an oxidized and reduced form, abbreviated as NAD^+^ and NADH respectively. NAD is a coenzyme central to metabolism, mainly involved in redox reactions. In addition, it is also participated in several post-translational modifications to regulate DNA damage response or gene expression, most notably as a substrate of the PARP enzyme family in ADP-Ribosylation and SIRT enzyme family in deacetylation. Three major biosynthetic pathways of NAD^+^, coordinated by a series of specialized synthesizing enzymes, have been adopted by different cell types and tissues^41–42^, including: quinolinate phosphoribosyltransferase (QAPRT)-mediated de novo synthesis pathway, nicotinate phosphoribosyltransferase (NAPRT)-mediated Preiss-Handler (PH) synthesis pathway and NAMPT-mediated salvage pathway. Because of the vital functions of NAD^+^, the enzymes involved in NAD metabolism are favorable targets for drug discovery.

NAMPT is a rate-limiting enzyme that catalyzes NAD^+^ synthesis in the salvage pathway.^43^ By combining whole-genome CRISPR screening and pan-cancer genetic dependency mapping, Nakada and colleagues have identified NAMPT as AML dependencies governing NAD^+^ biosynthesis.^44^ Since 1997, scientists have designed a panel of NAMPT inhibitors, including FK866, CHS-828, GMX1777, KPT-9274. Although very potent *in vitro*, they all face certain challenges during clinical investigation. Take FK866 as an example: (1) It shows great inhibition of the enzymatic activity of NAMPT initially, but cells respond quickly by synthesis more proteins, leading to high demand for the inhibitor dosage. This phenomenon was also recapitulated in our experimental setting (Fig. 5E). (2) Traditional enzymatic inhibitor like FK866 is incapable to antagonize the pro-tumorigenic function of eNAMPT. (3) Notably, FK866 shows severe thrombocytopenia side effects.^45^ Therefore, there is an urgent need to either improve existing inhibitors or develop brand-new inhibitors with a novel mechanism of action (MOA).

To our knowledge, this study is the first report to apply PROTAC technology to block the function of NAMPT by protein degradation. Our PROTAC compounds, **630120** and **630121**, couple the NAMPT inhibitor FK866 to the CRBN ligand lenalidomide. With the assistance of the compounds, CRBN is recruited to ubiquitinate NAMPT, the latter is then degraded by the proteasome. Interestingly, we found that NAMPT is a relatively stable protein, whose degradation under natural conditions is mainly through lysosome-dependent rather than proteasome-dependent pathway (Fig. 1). Following biochemical characterizations have confirmed the great sensitivity (Fig. 3), selectivity (Fig. 4A-B) and clear MOA (Fig. 4C-F) of these PROTAC compounds. Most importantly, biological characterizations with hematologic tumor cell line HL60 and MLOT4 as models have provided compelling evidence that both PROTAC compounds have equivalent inhibition on NAD^+^ production, but outperform FK866 in killing effect on these tumor cells (Fig. 5A-D). Mechanistically, PROTAC-mediated eNAMPT degradation and downstream Erk1/2 inactivation (Fig. 5E-F) may contribute significantly to this superior efficacy of the compounds.

PROTAC technology has developed rapidly in recent years. Its main advantages include the type of catalytic action, high selectivity, the potential to target traditionally undruggable proteins and the ability to overcome mutation-driven therapeutic resistance. However, this technology still has certain issues to be improved, such as poor oral absorption caused by large molecular weight and strong molecular rigidity.^34^ Our compounds, especially **630121**, overcame these shortcomings by rational design and showed very promising oral absorption and utilization in the pharmacokinetic assay in rats, which deserves more intensive pre-clinical research in the near future.

In summary, two NAMPT-targeting PROTAC compounds have been designed to achieve NAMPT degradation in a broad spectrum of tumor cell lines, thus interfering with the metabolic fitness of tumor cells. Our compounds could simultaneously eliminate both iNAMPT and eNAMPT proteins, and achieve a better killing effect than FK866 in multiple hematological tumor cell models.

## Acknowledgments

The authors thank the Multi-Omics Core Facility (MOCF) at the School of Life Science and experiments. The authors also thank the Molecular and Cell Biology Core Facility (MCBCF) at the School of Life Science and Technology, ShanghaiTech University for technical support with flow cytometry. This work was supported by the Ministry of Science and Technology of China (2018YFC1004603 to GF), the National Natural Science Foundation of China (32070776 and 31872831 to GF), the Shanghai Science and Technology Commission (19JC1413800 to GF), the Shanghai Shuguang program (19SG55 to GF) and a ShanghaiTech University startup grant.

## Author contribution

XZ, XY and GF designed the study. XZ, HL and LC performed the experiments; YC made the CRBN KO cell lines; XZ, HL, BJ, XY and GF interpreted the data; XZ, HL, LC and GF wrote the paper.

The authors declare that they have no competing interests.

## Data availability

The MS data have been deposited to the iProX Integrated Proteome Resources (https://www.iprox.cn/) with the dataset identifiers IPX0003508000. Other raw and analyzed data files are available from the corresponding author upon reasonable request.

