## supplementary data for "Addressing Enzymatic-Independent Tumor-Promoting Function of NAMPT via PROTAC-Mediated Degradation"

### Supporting Information

#### Pharmacokinetic Studies

The pharmacokinetic properties of compounds in rats were determined as follows. The compound was administered intravenously to the femoral vein of male Sprague Dawley rats as a solution in NMP/PEG200 (1:9, v/v) at 2 mg/kg, or as a 10 mg/kg NMP/PEG200 (1:9, v/v) solution for gavage. Serial plasma samples were collected by sublingual vein bleeding for 24 h (0.083, 0.25, 0.5, 1, 2, 4, 8, 24 h) from one group of rats (n = 3). Samples were extracted and the concentration of compound was determined by LC−MS/MS. LOQ was set to 2.5 ng/mL.

#### Chemistry General Procedure

All chemicals were obtained from commercial suppliers (Adamas and Alfa, bidepharm, ChemShuttle), and used without further purification, unless otherwise indicated. Flash chromatography was carried out on silica gel (200–300 mesh). Analytical TLC was performed on Haiyang ready-to-use plates with silica gel 60 ($F_{254}$). All new compounds were characterized by $^1$H NMR, $^{13}$C NMR, HRMS. $^1$H NMR,$^{13}$C NMR spectra were recorded on Bruker AVANCE III 500 MHZ (operating at 500 MHz for $^1$H NMR,$^{13}$C NMR), chemical shifts were reported in ppm relative to the residual $d_6$-DMSO (2.50 ppm $^1$H, 39.52 ppm $^{13}$C), $d_4$-Methanol(3.30 ppm $^1$H, 49 ppm $^{13}$C),and coupling constants ($J$) are given in Hz. Multiplicities of signals are described as follows: s --- singlet, br.--- broad singlet, d --- doublet, t --- triplet, m --- multiple. High Resolution Mass spectra were recorded on AB Triple 4600 spectrometer with acetonitrile and water as solvent. The final compounds were all purified by C18 reverse phase preparative HPLC column with solvent A (0.5% HCl in $H_2O$) and solvent B (MeCN) as eluents.

**(*E*)-N-(4-(piperidin-4-yl)butyl)-3-(pyridin-3-yl)acrylamide (3)**

The compound (*E*)-3-(pyridin-3-yl)acrylic acid (500 mg, 3.35 mmol) **1**,*tert*-butyl 4-(4-aminobutyl)piperidine-1-carboxylate(860 mg,3.35 mmol) **2**, HATU (1.5 g, 4.0 mmol) and DIPEA(1.3 g,10 mmol), DMF (10 mL) were added to a 50 mL single bottle. After being stirred overnight at 60 ℃, the crude mixture was cooled down to room temperature, diluted with 70% brine aqueous (50 mL), extracted with EtOAc. After that, the combined organic layer was washed with water and brine, then dried over $Na_2SO_4$ and concentrated in vacuum to afford crude oil. After solvent evaporation, the residue was purified by column chromatography on silica gel (5% $CH_3OH$ in DCM) to give the intermediate (880 mg, crude yield 68%) as yellow solid.

NMR spectra shows in Figures S3.

The obtained intermediate was then added to a 25 mL single bottle with TFA solution (4ml TFA in 10 mL DCM) and stirred at room temperature for 12 h. When the reaction was completed, the mixture was purified by preparative HPLC (10 to 90% acetonitrile/0.05% TFA in $H_2O$) to obtain the target compound (600 mg,92%) as yellow oily liquid. $^1$H NMR (500 MHz, Methanol-$d_4$) δ 9.12 (s, 1H), 8.85 - 8.80 (m, 2H), 8.12 – 7.98 (m, 1H), 7.63 (d, *J* = 15.8 Hz, 1H), 7.02 (d, *J* = 15.9 Hz, 1H), 3.44 – 3.27 (m, 5H), 3.05 - 2.95 (m, 2H), 1.95 (dd, *J* = 13.6, 3.4 Hz, 2H), 1.60 (p, *J* = 7.1 Hz, 3H), 1.45 – 1.27 (m, 7H). $^{13}$C NMR (126 MHz, Methanol-$d_4$) δ 166.63, 144.34, 143.27, 142.88, 136.57, 134.02, 129.05, 128.60, 45.29, 40.56, 36.60, 34.72, 30.31, 29.95, 24.73, 18.74, 17.30. MS(ESI) m/z: 288.2074 $[M+H]^+$.

**(*E*)-N-(4-(1-(4-(piperazin-1-yl)benzoyl)piperidin-4-yl)butyl)-3-(pyridin-3-yl)acrylamide (5)**

The title compound (yellow oily liquid, 600mg, 60% yield over two steps) was synthesized according to procedures for the preparation of **3**. $^1$H NMR (500 MHz, Methanol-$d_4$) δ 9.13 (s, 1H), 8.87 (dd, *J* = 18.8, 6.9 Hz, 2H), 8.15 (dd, *J* = 8.2, 5.7 Hz, 1H), 7.64 (d, *J* = 15.8 Hz, 1H), 7.50 - 7.42 (m, 2H), 7.26 - 7.16 (m, 2H), 7.05 (d, *J* = 15.8 Hz, 1H), 4.98 (br, 4H), 3.63 (s, 4H), 3.43 (s, 4H), 3.34 (t, *J* = 7.0 Hz, 2H), 1.94 – 1.75 (m, 2H), 1.70 - 1.56 (m, 3H), 1.49

- 1.40 (m, 2H), 1.40 – 1.32 (m, 2H), 1.31 – 1.19 (m, 2H). $^{13}C$ NMR (126 MHz, Methanol-$d_4$) δ 170.98, 165.15, 151.30, 144.11, 141.01, 140.71, 135.51, 132.25, 128.93, 128.19, 127.49, 115.81, 45.55, 42.96, 39.30, 35.39 (d, $J$ = 10.0 Hz), 29.05, 23.64. MS(ESI) m/z: 476.3024 $[M+H]^+$.

NMR spectra shows in Figures S4.

**7-(2-(2,6-dioxopiperidin-3-yl)-1-oxoisoindolin-4-yl)hept-6-yn-1-yl methanesulfonate (7)**

The compound 3-(4-bromo-1-oxoisoindolin-2-yl)piperidine-2,6-dione (800 mg,2.5 mmol) **6** , hept-6-yn-1-ol (560 mg, 5mmol), $Pd(PPh_3)_2Cl_2$ (175mg,0.25 mmol), CuI (95 mg,0.5 mmol) and anhydrous DMF (5 mL) were added to a 50 mL single bottle and stirred 5 min at room temperature under Ar atmosphere. After the mixture was added 2.5 mL anhydrous $Et_3N$, the reaction was stirred at 80 °C for 12 h, Then the crude mixture was cooled down to room temperature, diluted with $H_2O$ (50 mL), extracted with EtOAc. After that, the combined organic layer was washed with water and brine, then dried over $Na_2SO_4$ and concentrated in vacuum to afford crude oil. After solvent evaporation, the residue was purified by column chromatography on silica gel (5% $CH_3OH$ in DCM) to give the compound (800 mg, crude yield 90%) as white solid.

The obtained intermediate dissolved in 30 mL DCM was added to a 25 mL single bottle with $Et_3N$ (676 mg, 6.7 mmol) and stirred at 0 ℃ for 15 min. When the reaction was completed, the mixture was purified by column chromatography on silica gel (5% $CH_3OH$ in DCM) to obtain target compound (887mg, 91%) as white solid. $^1H$ NMR (500 MHz, DMSO-$d_6$) δ 10.99 (s, 1H), 7.71 (dd, $J$ = 7.5, 1.1 Hz, 1H), 7.64 (dd, $J$ = 7.6, 1.1 Hz, 1H), 7.52 (t, $J$ = 7.6 Hz, 1H), 5.13 (dd, $J$ = 13.3, 5.1 Hz, 1H), 4.45 (d, $J$ = 17.6 Hz, 1H), 4.31 (d, $J$ = 17.6 Hz, 1H), 4.22 (t, $J$ = 6.4 Hz, 2H), 3.15 (s, 3H), 2.96 - 2.86 (m, 1H), 2.64 – 2.55 (m, 1H), 2.52 (br, 2H), 2.47 - 2.40 (m, 1H), 2.06 - 1.97 (m, 1H), 1.81 – 1.66 (m, 2H), 1.62 (p, $J$ = 7.0 Hz, 2H), 1.55 - 1.47 (m, 2H). $^{13}C$ NMR (126 MHz, DMSO-$d_6$) δ 172.88, 170.99, 167.67, 143.74, 134.10, 131.99, 128.60, 122.63, 118.80, 96.12, 76.53, 70.40, 51.65, 46.97, 36.55, 31.20, 28.02, 27.46, 24.29, 22.34, 18.64. MS(ESI) m/z: 433 $[M+H]^+$.

NMR spectra shows in Figures S5.

**8-(2-(2,6-dioxopiperidin-3-yl)-1-oxoisoindolin-4-yl)oct-7-yn-1-yl methanesulfonate (8)**

The title compound (white solid, 914mg, 82% yield over two steps) was synthesized according to procedures for the preparation of **7**. $^{1}$H NMR (500 MHz, DMSO-$d_6$) δ 11.00 (s, 1H), 7.70 (dd, *J* = 7.6, 1.1 Hz, 1H), 7.63 (dd, *J* = 7.6, 1.1 Hz, 1H), 7.51 (t, *J* = 7.6 Hz, 1H), 5.14 (dd, *J* = 13.3, 5.1 Hz, 1H), 4.45 (d, *J* = 17.6 Hz, 1H), 4.31 (d, *J* = 17.7 Hz, 1H), 4.20 (t, *J* = 6.4 Hz, 2H), 3.15 (s, 3H), 2.95 - 2.86 (m, 1H), 2.64 - 2.56 (m, 1H), 2.48 (d, *J* = 6.9 Hz, 2H), 2.47 – 2.41 (m, 1H), 2.04 - 1.97 (m, 1H), 1.73 – 1.64 (m, 2H), 1.59 (p, *J* = 7.0 Hz, 2H), 1.51 – 1.35 (m, 4H). $^{13}$C NMR (126 MHz, DMSO-$d_6$) δ 172.89, 171.00, 167.67, 143.74, 134.08, 131.97, 128.60, 122.61, 118.84, 96.27, 76.44, 70.41, 51.64, 46.96, 36.54, 31.20, 28.43, 27.81 (d, *J* = 21.7 Hz), 24.43, 22.37, 18.67. MS(ESI) m/z: 447 $[M+H]^+$.

NMR spectra shows in Figures S6.

**(*E*)-N-(4-(1-(4-(4-(7-(2-(2,6-dioxopiperidin-3-yl)-1-oxoisoindolin-4-yl)hept-6-yn-1-yl)piperazin-1-yl)benzoyl)piperidin-4-yl)butyl)-3-(pyridin-3-yl)acrylamide (SIAIS630120)**

The compound (*E*)-N-(4-(1-(4-(piperazin-1-yl)benzoyl)piperidin-4-yl)butyl)-3-(pyridin-3-yl)acrylamide (20 mg, 0.04 mmol) **5** ,7-(2-(2,6-dioxopiperidin-3-yl)-1-oxoisoindolin-4-yl)hept-6-yn-1-yl methanesulfonate (20.7mg, 0.048mmol) **7** , DIPEA(15.6 mg,0.12 mmol), NaI(17.8 mg,0.12 mmol)and DMF (2 mL) were added to a 25 mL single bottle. After being stirred 6 h at 60 ℃, the resulting mixture was purified by preparative HPLC (10 to 90% acetonitrile/0.05% HCl in $H_2O$) to obtain target compound (5.8mg. 25%) as white solid.$^{1}$H NMR (500 MHz, DMSO-d6) δ 11.00 (s, 1H), 8.89 (s, 1H), 8.70 – 8.66 (m, 1H), 8.28 - 8.25 (m, 2H), 7.71 (d, *J* = 7.5 Hz, 1H), 7.65 (d, *J* = 7.5 Hz, 1H), 7.56 – 7.45 (m, 2H), 7.28 (d, *J* = 8.5 Hz, 2H), 7.01 (d, *J* = 8.5 Hz, 2H), 6.82 (d, *J* = 15.9 Hz, 2H), 5.16 (dd, *J* = 13.3, 5.1 Hz, 1H), 4.47 (d, J = 17.7 Hz, 1H), 4.33 (d, *J* = 17.7 Hz, 1H), 3.89 (d, *J* = 12.9 Hz, 2H),3.55 (br, 4H), 3.24 – 3.16 (m, 5H), 3.14 – 3.04 (m, 5H), 2.98 - 2.88 (m, 1H), 2.65 – 2.56 (m, 1H), 2.55 – 2.51 (m, 2H), 2.49 - 2.43 (m, 1H), 2.07 - 1.97 (m, 1H), 1.84 - 1.75 (m, 2H), 1.69 -

1.58 (m, 4H), 1.53 - 1.42 (m, 5H), 1.35 - 1.29 (m, 2H), 1.26 - 1.22 (m, 3H), 1.12 – 0.97 (m, 2H).$^{13}$C NMR (126 MHz, DMSO-d6) δ 172.91, 171.03, 168.96, 167.64, 164.08, 150.17, 143.77, 134.09, 133.84, 131.99, 128.63, 128.44, 127.03, 125.94, 125.27, 122.68, 118.76, 114.79, 96.00, 76.62, 55.16, 51.64, 50.39, 47.00, 44.69, 38.70, 35.53, 35.41, 31.20, 29.24, 27.43, 25.28, 23.49, 22.43 (d, *J* = 9.8 Hz), 18.49. MS(ESI) m/z: 812.4496 [M+H]$^+$.

NMR spectra shows in Figures S7.

**(*E*)-N-(4-(1-(4-(4-(8-(2-(2,6-dioxopiperidin-3-yl)-1-oxoisoindolin-4-yl)oct-7-yn-1-yl)piperazin-1-yl)benzoyl)piperidin-4-yl)butyl)-3-(pyridin-3-yl)acrylamide (SIAIS630121)**

The title compound (white solid, 5.7mg, 24% yield) was synthesized according to procedures for the preparation of **SIAIS630120**. $^1$H NMR (500 MHz, Methanol-d4) δ 8.84 (d, J = 2.2 Hz, 1H), 8.63 (dd, J = 5.2, 1.5 Hz, 1H), 8.37 - 8.32 (m, 1H), 7.75 (d, J = 6.6 Hz, 1H), 7.71 (dd, J = 8.1, 5.2 Hz, 1H), 7.63 – 7.55 (m, 2H), 7.51 (t, J = 7.6 Hz, 1H), 7.36 (d, J = 8.7 Hz, 2H), 7.07 (d, J = 8.9 Hz, 2H), 6.81 (d, J = 15.9 Hz, 1H), 5.19 (dd, J = 13.3, 5.2 Hz, 1H), 4.53 (d, J = 17.4 Hz, 1H), 4.47 (d, J = 17.4 Hz, 1H), 3.95 (br, 1H), 3.67 (br, 2H), 3.39 - 3.32 (m, 6H), 3.25 – 3.19 (m, 4H), 3.18 – 3.11 (m, 2H), 2.95 - 2.86 (m, 1H), 2.84 - 2.75 (m, 1H), 2.57 – 2.49 (m, 3H), 2.24 - 2.15 (m, 1H), 1.86 - 1.80 (m, 2H), 1.70 (p, J = 6.9 Hz, 3H), 1.65 – 1.55 (m, 5H), 1.53 – 1.39 (m, 4H), 1.37 – 1.27 (m, 3H), 1.24 - 1.07 (m, 2H). $^{13}$C NMR (126 MHz, Methanol-$d_4$) δ 174.69, 172.32 (d, *J* = 11.1 Hz), 171.01, 167.28, 152.17, 147.08, 145.28, 139.35, 136.11, 135.78, 132.92, 129.69 (d, *J* = 4.6 Hz), 128.84, 126.69, 126.40, 123.76, 120.93, 116.85, 97.12, 77.40, 58.00, 53.70, 52.85, 47.10, 40.58, 37.11 (d, *J* = 10.7 Hz), 32.35, 30.53, 29.39 (d, *J* = 3.7 Hz), 27.12, 24.99 (d, *J* = 3.4 Hz), 24.11, 19.94. MS(ESI) m/z: 826.4655 [M+H]$^+$.

NMR spectra shows in Figures S8.

**Figure Legends**

**Figure S1. Construction and verification of NAMPT knock-in HEK293T cell line using Tild-CRISPR.**

(A) Genomic DNA sequence map from Snap Gene software of 293FT KI cell line. (B)

Different fields of fluorescence microscope imaging. Magnification: 10 X, ruler: 100 μm.

**Figure S2. Drug metabolism and pharmacokinetic/pharmacodynamic profile of 630120 and 630121.**

Plasma concentration-time profiles of (A) **630120** and (B) **630121** in rats.

**Figure S3. $^{1}H$ and $^{13}C$ NMR spectra for compound 3**

**Figure S4. $^{1}H$ and $^{13}C$ NMR spectra for compound 5**

**Figure S5. $^{1}H$ and $^{13}C$ NMR spectra for compound 7**

**Figure S6. $^{1}H$ and $^{13}C$ NMR spectra for compound 8**

**Figure S7. $^{1}H$ and $^{13}C$ NMR spectra for compound 630120**

**Figure S8. $^{1}H$ and $^{13}C$ NMR spectra for compound 630121**

**A**

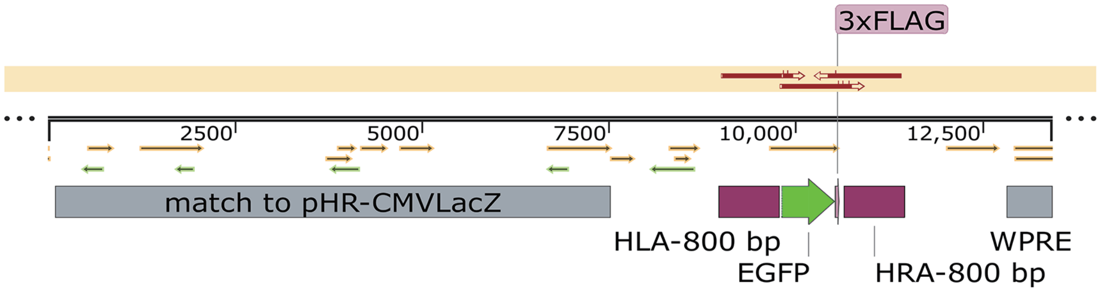


**B**

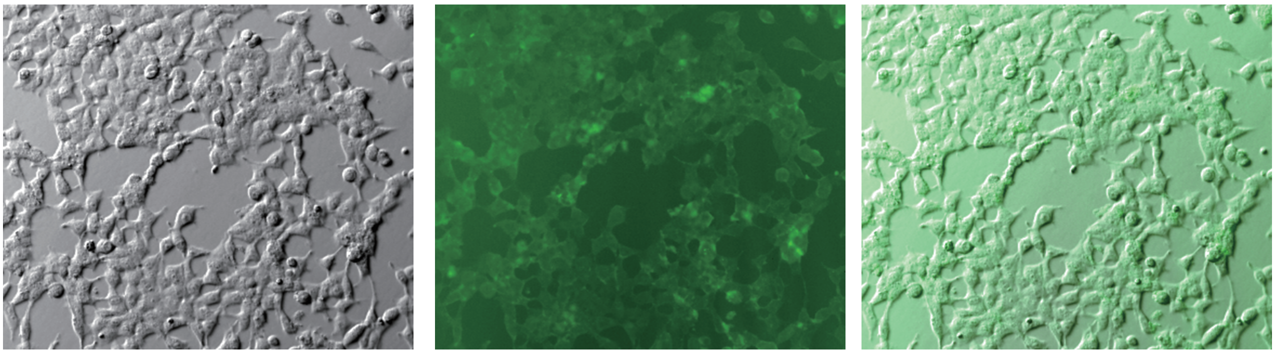

**Bright field** **FITC field** **Merged field**

A

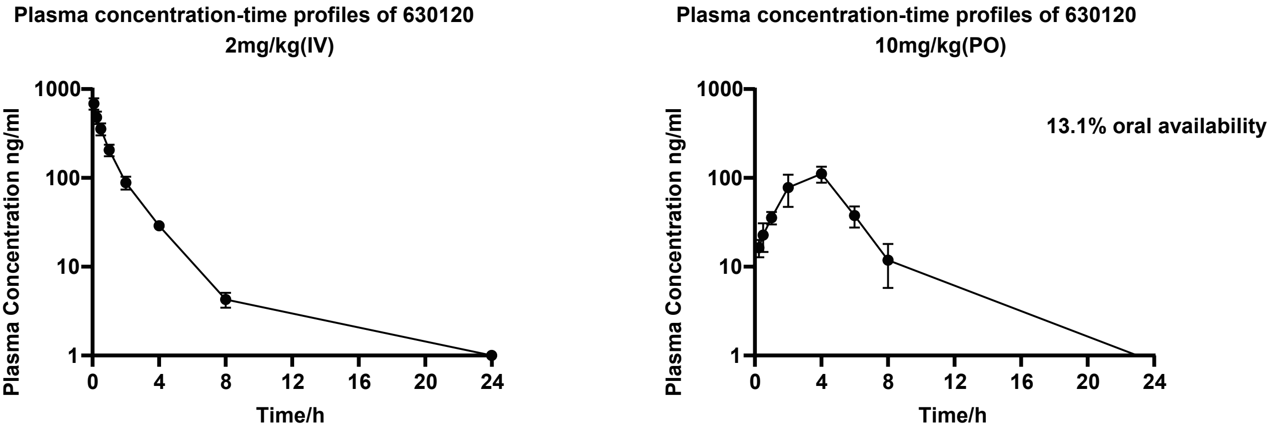
Plasma concentration-time profiles of 630120
2mg/kg(IV)
Plasma concentration-time profiles of 630120
10mg/kg(PO)
13.1% oral availability
Plasma Concentration ng/ml
Time/h
1000
100
10
1
0
4
8
12
16
20
24

B

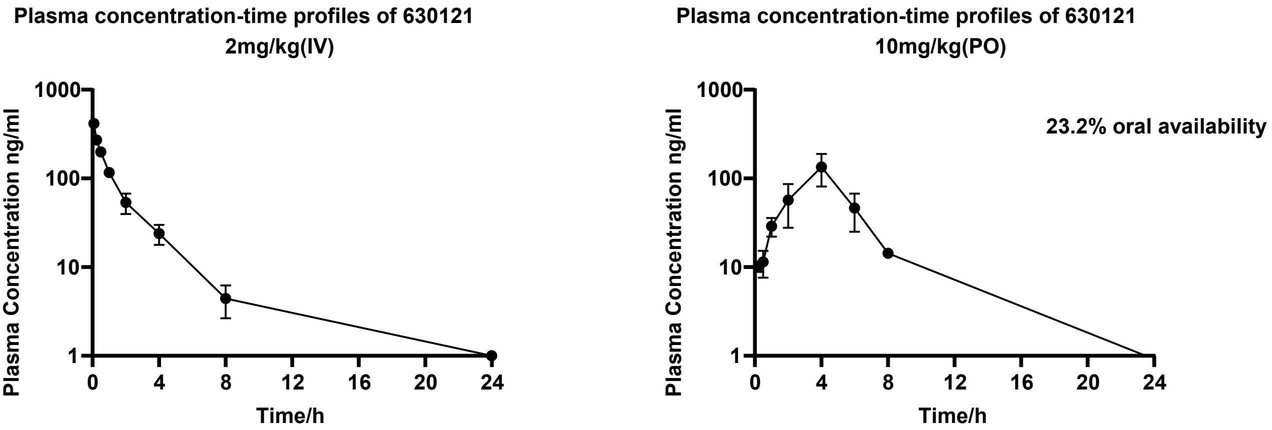
Plasma concentration-time profiles of 630121
2mg/kg(IV)
Plasma concentration-time profiles of 630121
10mg/kg(PO)
23.2% oral availability
Plasma Concentration ng/ml
Time/h
1000
100
10
1
0
4
8
12
16
20
24

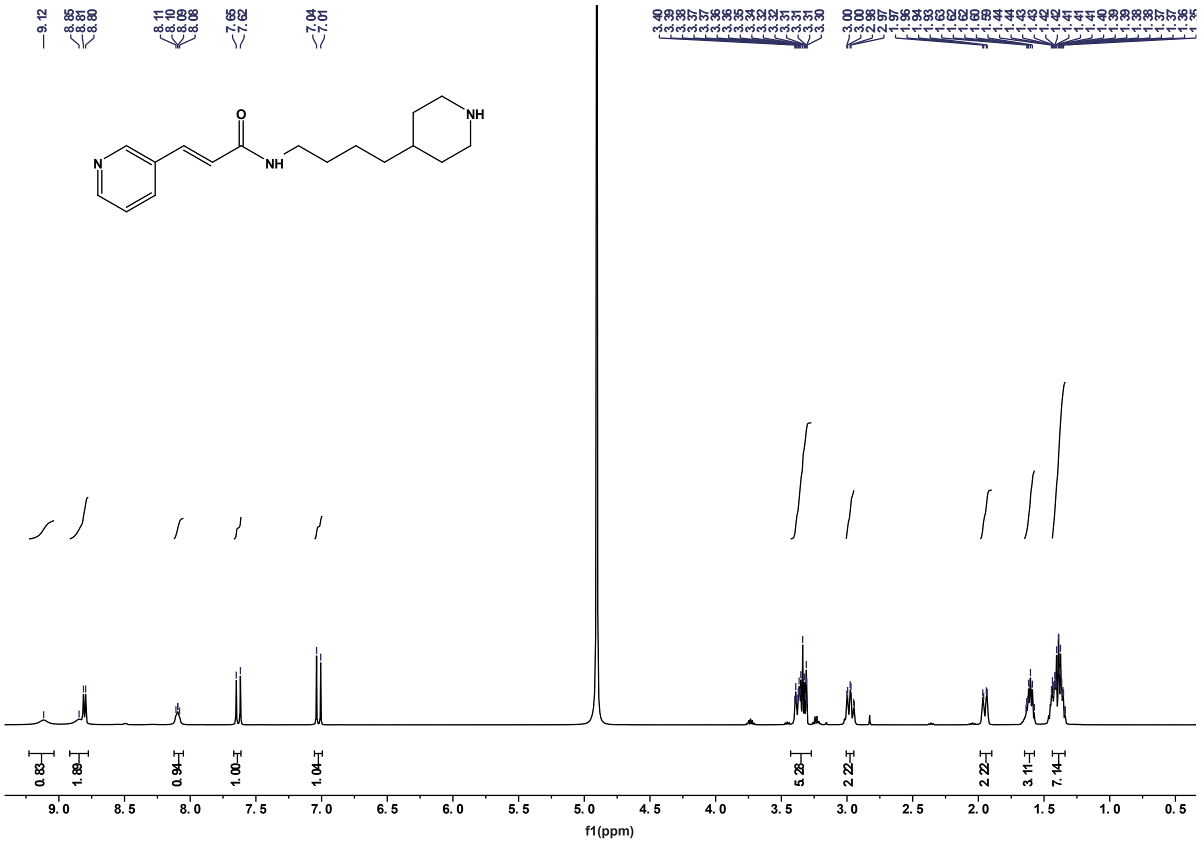
f1(ppm)

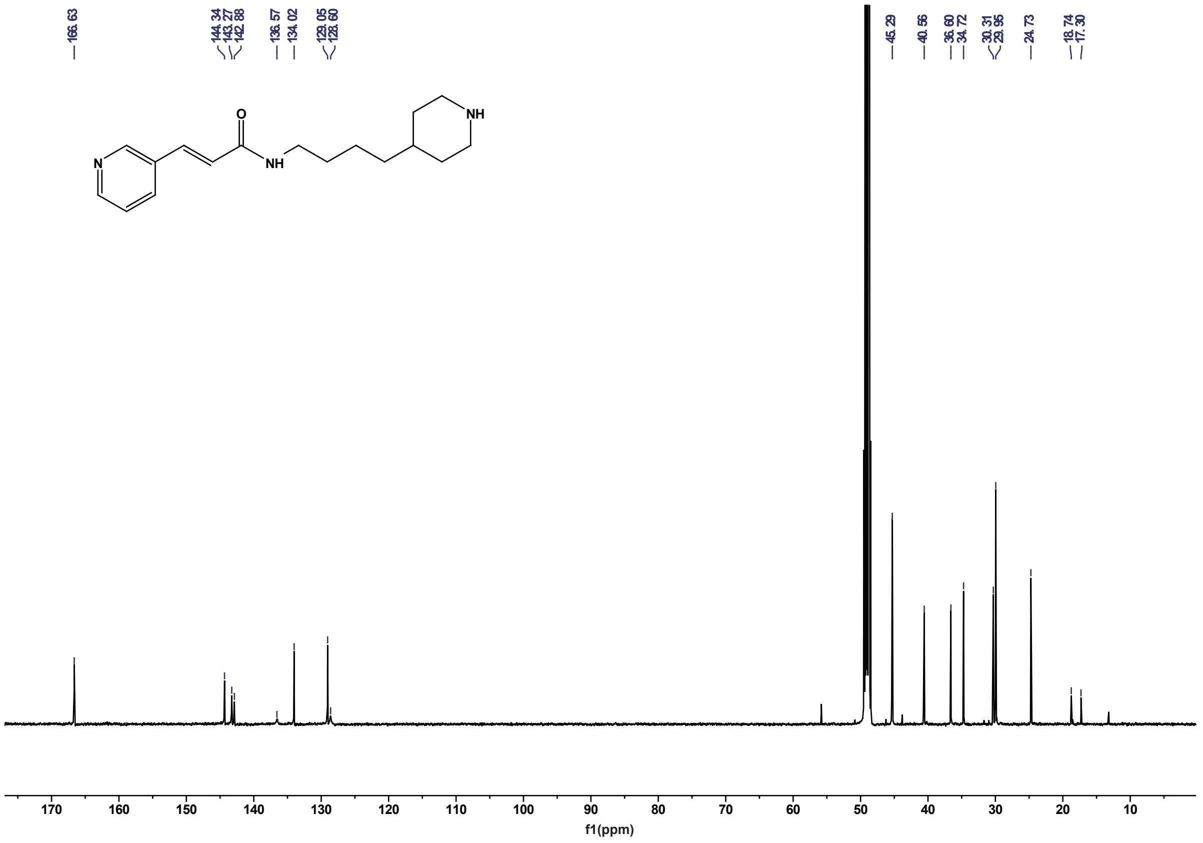
f1(ppm)

#### NAMPT PROTAC Fig S4

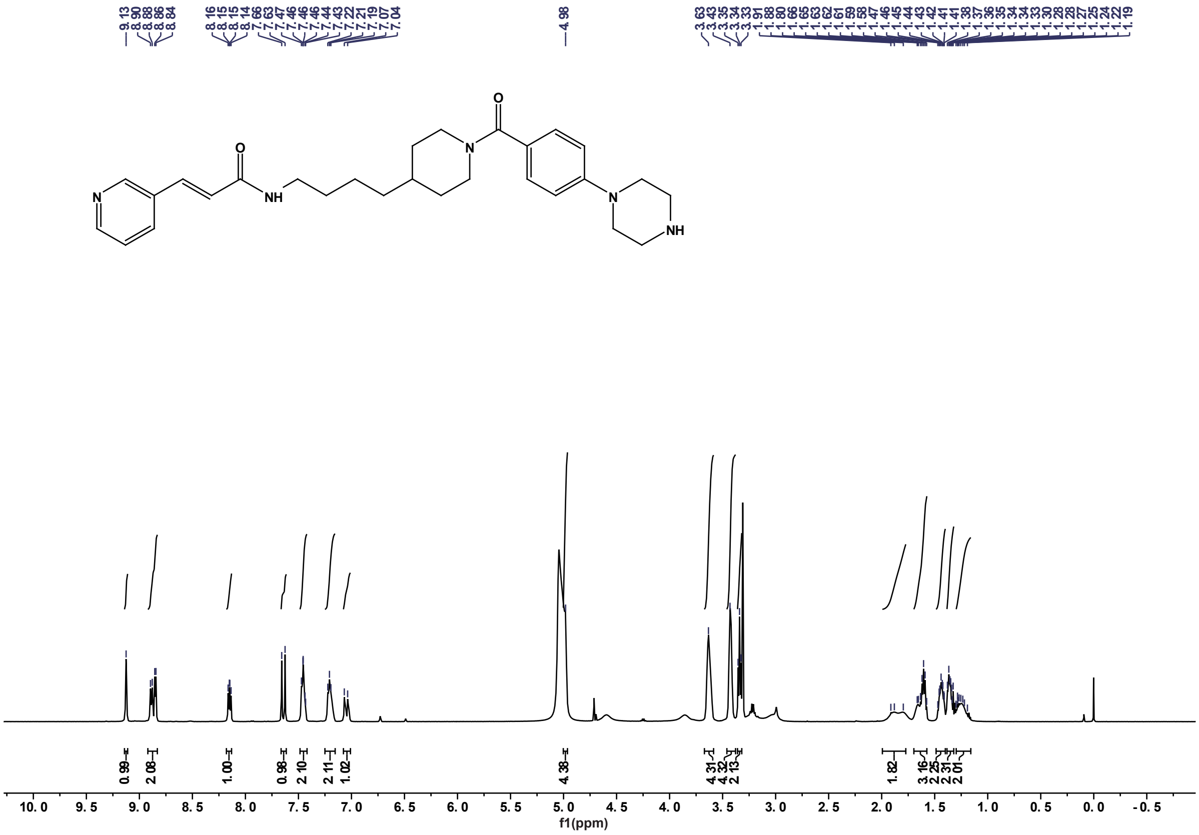


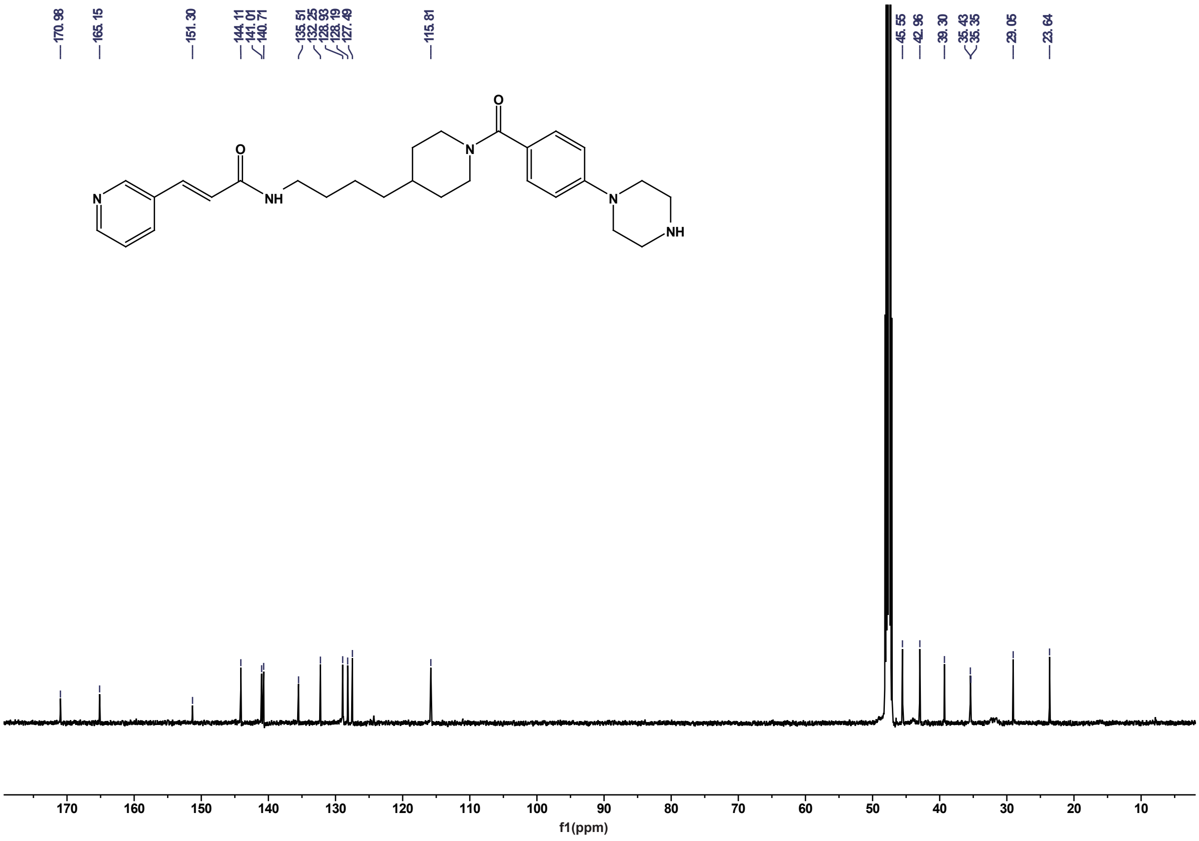

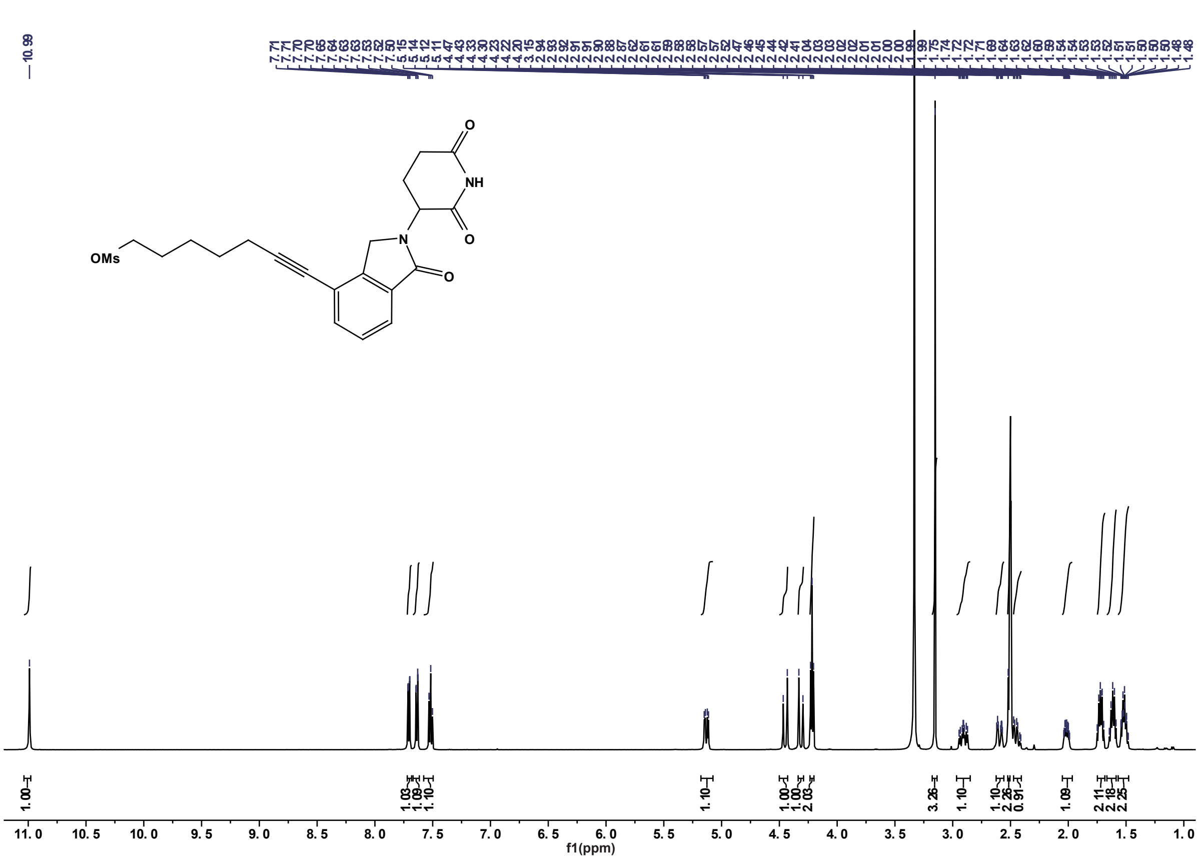
10.99
OMs
NH
O
N
f1(ppm)

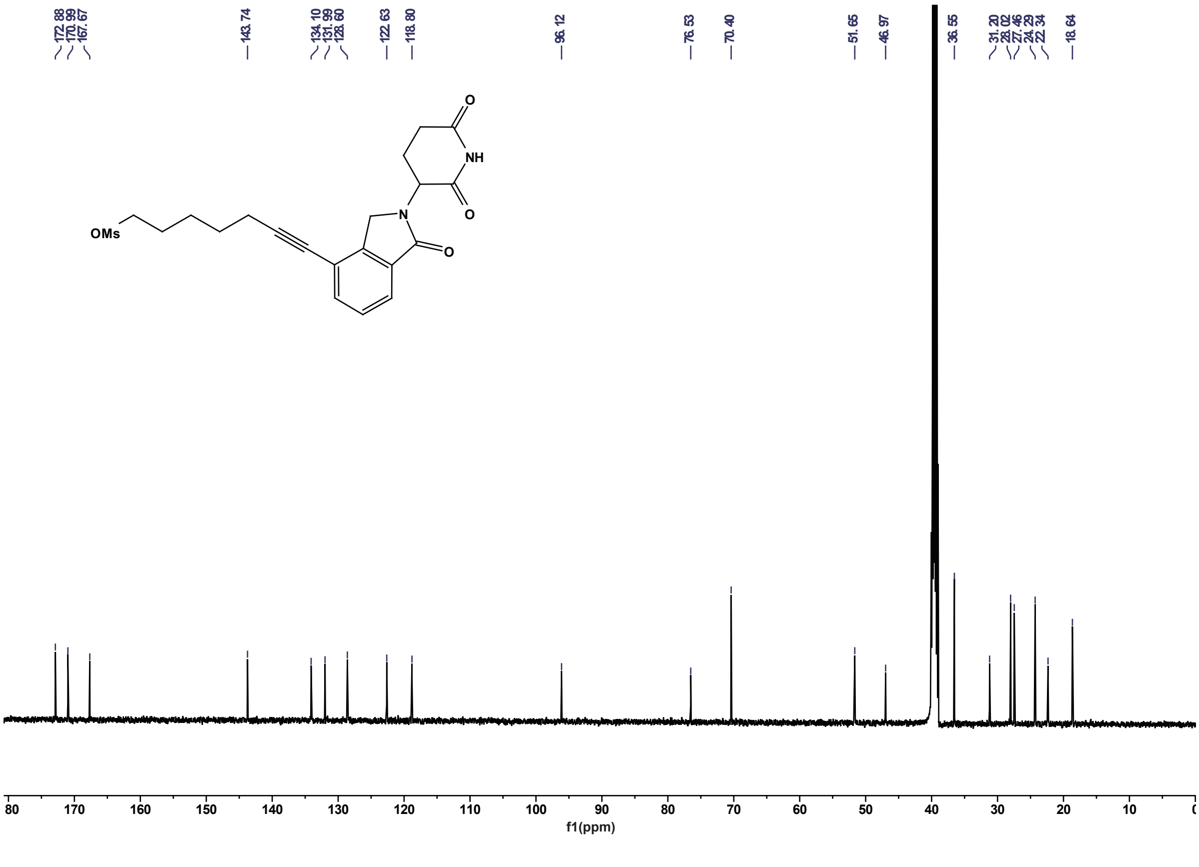
172.88
170.99
167.67
143.74
134.10
131.99
128.60
122.63
118.80
96.12
76.53
70.40
51.65
46.97
36.55
31.20
28.02
27.46
24.29
22.34
18.64
OMs
NH
O
N
f1(ppm)

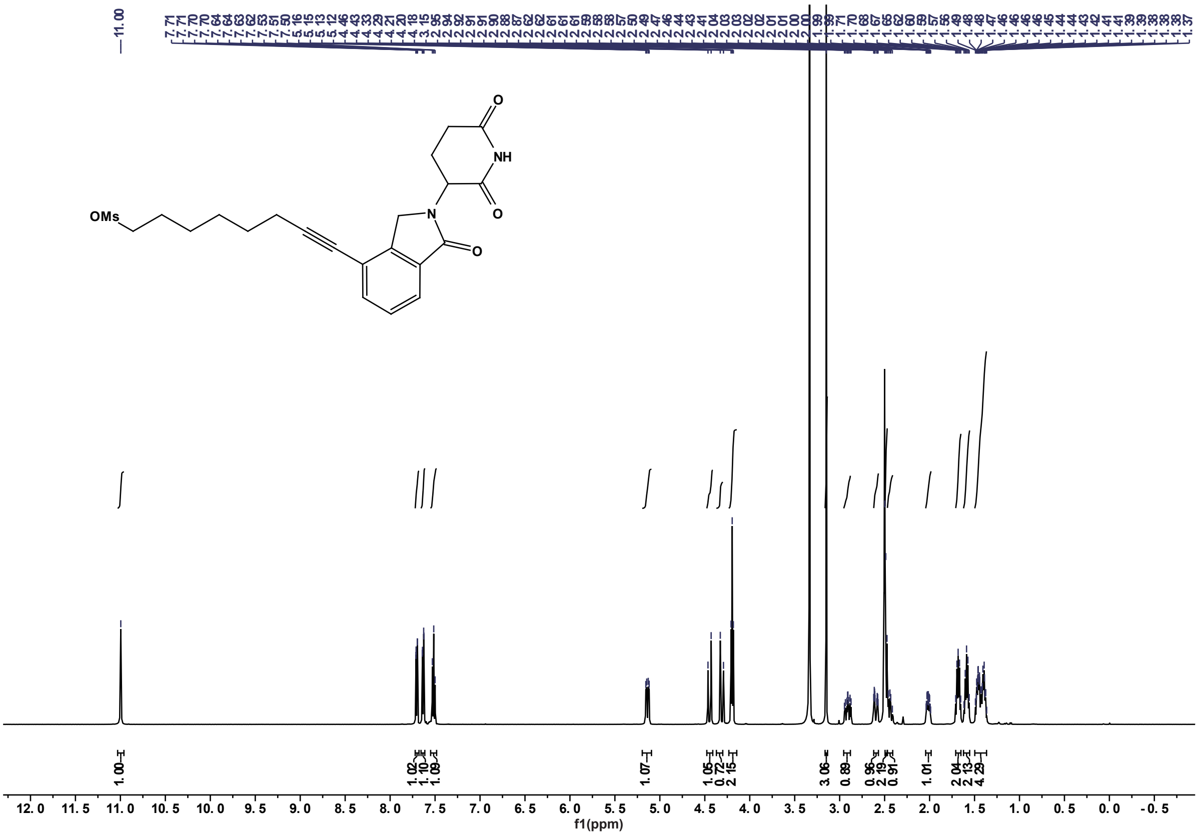

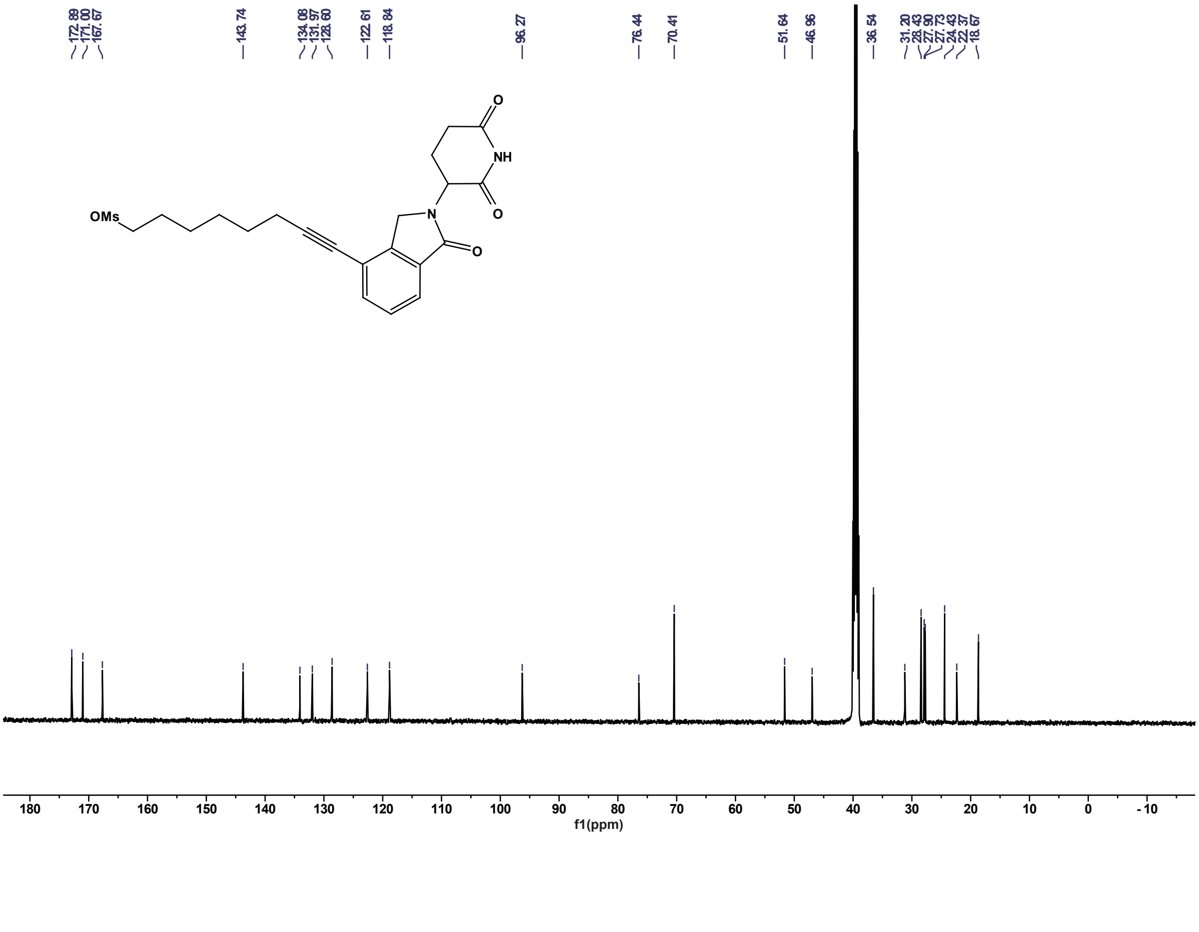

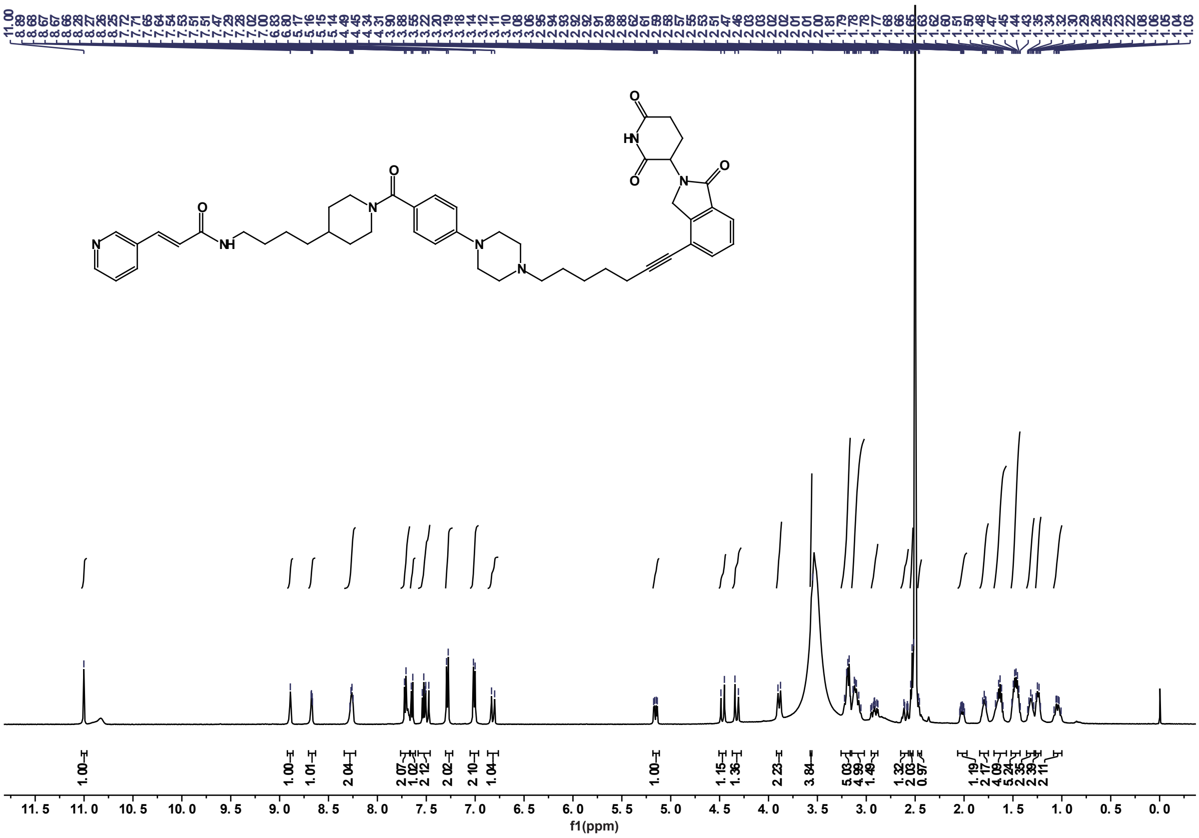
O
HN
O
N
O
O
N
O
N
H
N
N
f1(ppm)

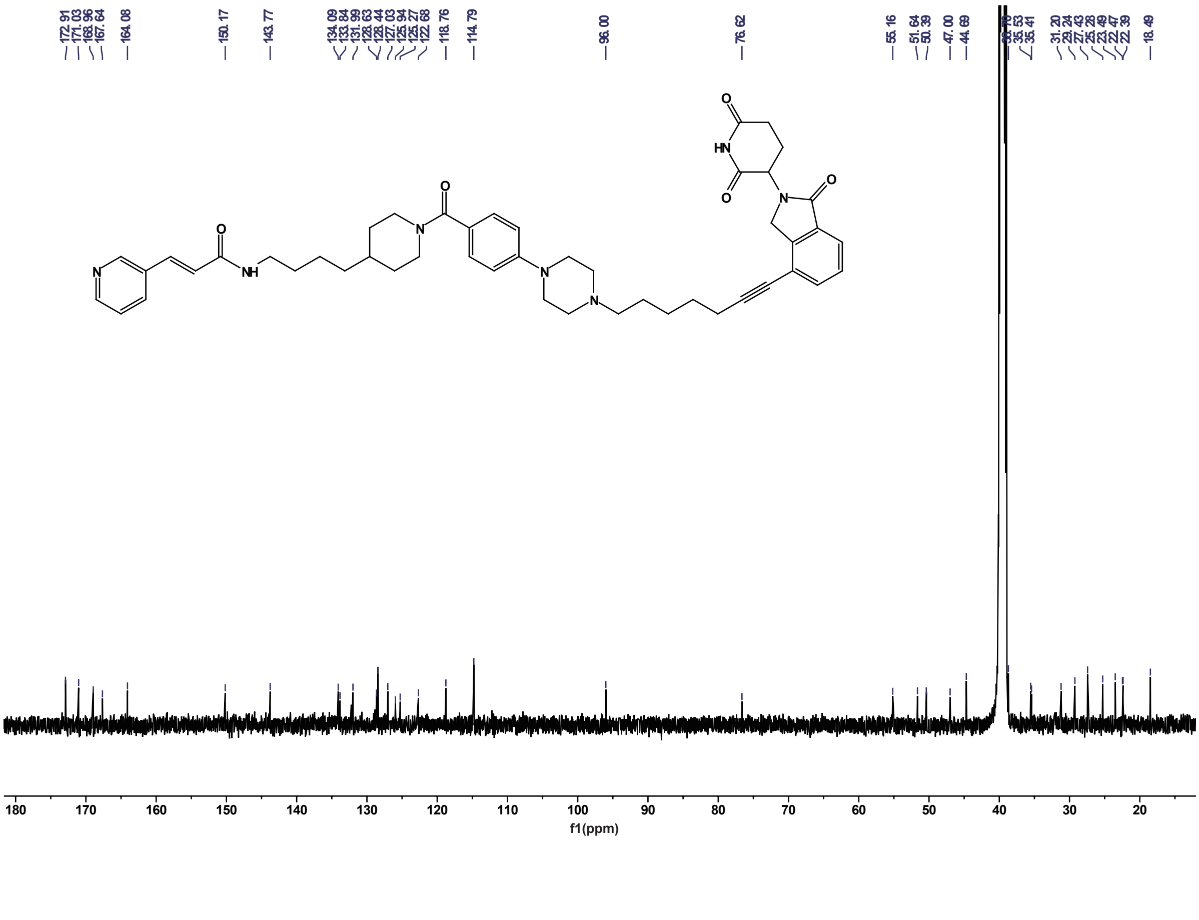
O
HN
O
N
O
O
N
O
N
H
N
N
f1(ppm)

#### NAMPT PROTAC Fig S8

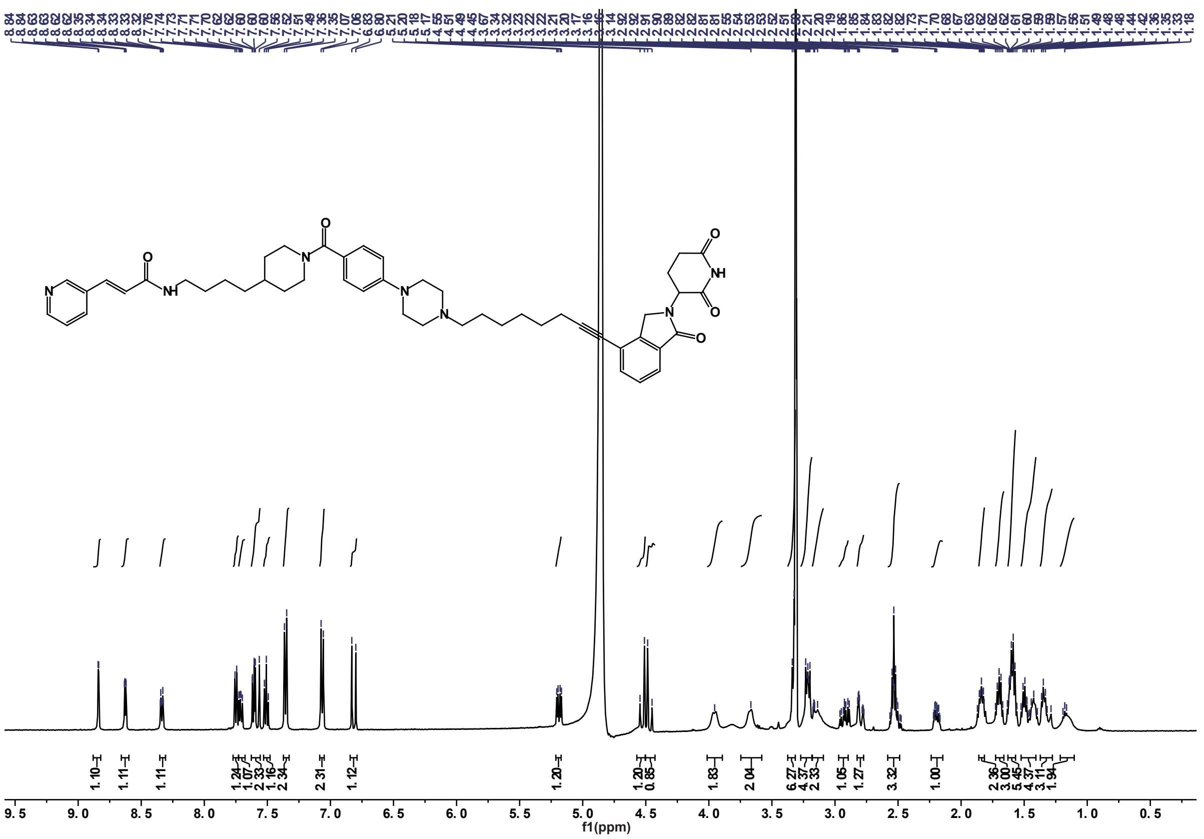

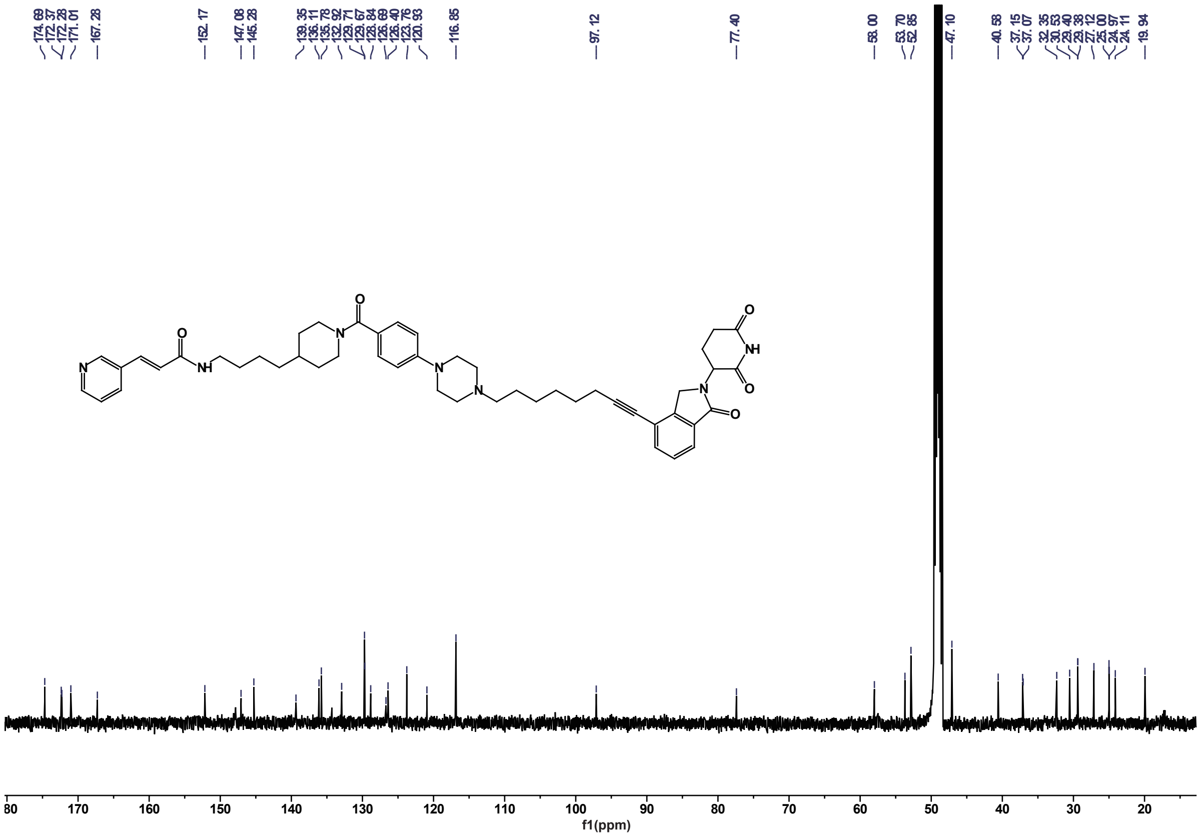
